# Prioritized experience replays on a hippocampal predictive map for learning

**DOI:** 10.1101/2020.03.23.002964

**Authors:** Hideyoshi Igata, Yuji Ikegaya, Takuya Sasaki

## Abstract

Hippocampal cells are central to spatial and predictive representations, and experience replays by place cells are crucial for learning and memory. Nonetheless, how hippocampal replay patterns dynamically change during the learning process remains to be largely elucidated. We found that when rats updated their behavioral strategies in response to a novel salient location, place cells increased theta sequences and sharp wave ripple-associated synchronous spikes that preferentially replayed salient locations and reward-related contexts in reverse order. The directionality and contents of the replays progressively varied with learning, including an optimized path that had never been exploited by the animals. Realtime suppression of sharp wave ripples during learning inhibited the rapid stabilization of optimized behavior. Our results suggest that hippocampal replays prioritize salient experiences and support the reinforcement of new behavioral policies.

**One-Sentence Summary:** Neuronal network mechanisms to internally rehearse learned information for updating behavioral strategy are revealed

## Main text

To adapt to continuous changes in the external environment, animals need to encode new salient episodes and update their behavioral policy. To achieve flexible spatial navigation in particular, hippocampal place cells are thought to provide fundamental neural spatial coding and constitute a cognitive map (*1*). This view has been proposed to extend to a predictive map theory in which place cells potentially encode expectations about animal’s future states (*2*). In addition to map representations, place cells that encode animals’ past or future trajectories are sequentially activated during sharp wave ripples (SWRs) within a short time window (~100 ms) in a phenomenon known as place cell replays (*3–5*), which are potentially appropriate to induce plasticity in the hippocampal circuit. A growing amount of evidence demonstrates that the frequency of SWR-associated place cell replays (or reactivation) is increased during learning of new, unexpected experiences (online replays) (*6*) and during rest/sleep states after learning of new experiences (offline replays) (*7*). Moreover, selective disruption of waking SWRs during spatial learning has been demonstrated to reduce subsequent task performance (*8*) and impair the stabilization and refinement of place cell maps (*9*). Taken together, these results suggest an essential role of SWR-associated replays in novel spatial learning.

Moreover, experience replays by place cells are suggested to be a key neuronal basis for a reinforcement learning framework (*10, 11*). When agents encounter a prediction error, such as a change in reward, experience replays may be an efficient mechanism for the evaluation of their experienced action-outcome associations by providing a solution to the temporal credit assignment problem, leading to updates of their behavioral policy to maximize future reward (*12, 13*). In line with this idea, empirical observations have demonstrated that the receipt of reward or novel experiences leads to increased rates of SWRs and coordinated reactivation of place cells (*6, 14*), and increased reward leads to increased reverse replays (*4, 15*).

Despite accumulating evidence and theory, the field still lacks key insights into how the contents of place cell replays progressively change to (re-)assign the values of salient locations as animals develop new navigation strategies in response to environmental changes. Theoretical works demonstrate that incorporating replay algorithms with prioritized memory access for learned salient locations (*16*), rather than random access from all stored memory, into a deep neural network—a strategy termed prioritized experience replays (*17*)—improves integrative learning in artificial agents. Whether living neuronal networks adopt the same computations to enhance learning capability remains an open question.

To address this issue, we designed a new spatial task in which rats ran from a starting area to checkpoint C_1_ (path S-C_1_), where they received a small (20 μl chocolate milk) reward, and then ran from C_1_ to a goal area (path C_1_-G), where they received a large (200 μl chocolate milk) reward (Fig. 1A and Fig. S1A-D). On a recording day, after the animals were well trained on this task, the rats first performed the same task, termed the prelearning phase. After 11-14 trials, the reward was replaced to a new checkpoint (C_2_) (Fig. 1A, right), requiring the rats to learn new trajectories through trial and error. After the replacement, the rats first persisted in traversing path S-C_1_-G; however, the reward was no longer presented. The rats then visited C_2_ via path G-C_2_. The rats gradually spent less time on path S-C_1_-G and instead settled on the most efficient trajectory, path S-C_2_-G (Fig. 1B-E and fig. S2). For each rat, a learning point was determined based on a moving-average learning curve, and the periods before and after the learning point were termed the learning phase and the postlearning phase, respectively. The rats required 26 ± 9 trials (approximately 30 min) to progress from reward replacement to the learning point. In total, paths S-C_1_, C_1_-G, G-C_2_, S-C_2_, and C_2_-G accounted for 93.9% of all animal trajectories (Fig. 1F and 1G). Compared with previous works, the concepts of this task were that (1) the consistency of task demands, irrespective of the locations of check points, enabled us to extract spatial and context-dependent neuronal codes (*18, 19*), and (2) an obvious learning point and the animals’ learning-dependent trajectories enabled us to analyze detailed hippocampal replay patterns associated with the rats’ internal evaluations of behavioral strategies.

**Fig. 1.**
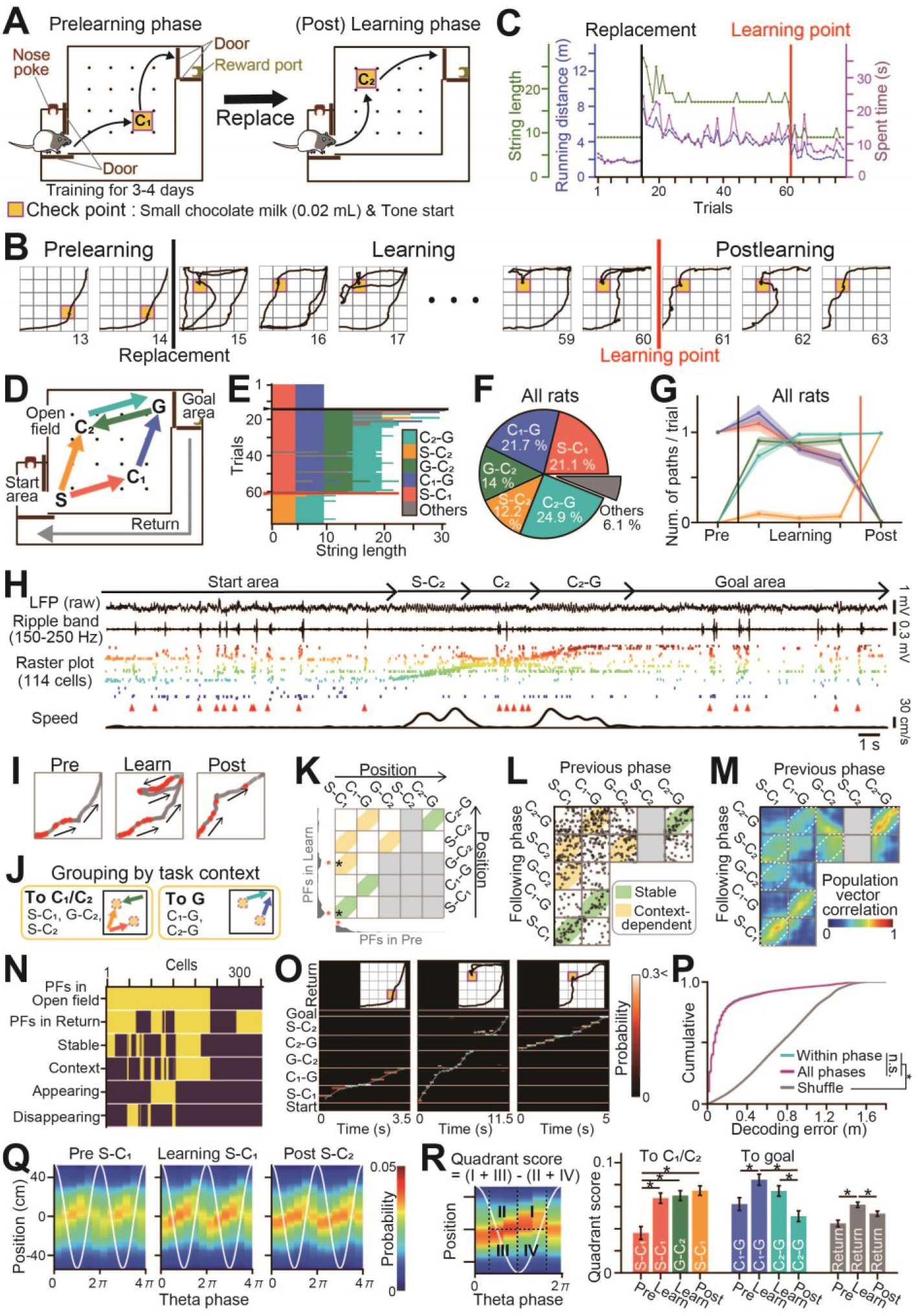
Behavioral performance and hippocampal place cell activity in the spatial learning task. (A) Overview of the task. The rat first took a trajectory from the start to the goal area via C_1_, where a reward was placed (path S-C_1_-G; prelearning phase). The areas were separated by automatic doors. The reward point was then changed to C_2_, and the rat learned to take a new trajectory via C_2_ (learning phase). (B) Representative trajectories observed from a single rat (all trajectories are shown in fig. S2A). The black and red vertical lines indicate reward replacement and a learning point defined based on a learning curve, respectively. Trial numbers are indicated below the images. (C) Changes in running distance, duration, and string length for the rat. (D) Five paths analyzed. (E) Changes in string length for the rat. The black and red horizontal lines indicate the replacement and learning points, respectively. (F) Proportions of paths taken by all five rats. (G) Changes in the number of paths per trial. The colors correspond to the paths in D. The thin shaded regions indicate the SEM. (H) (top to bottom) Original local field potential (LFP), ripple band (150-250 Hz)-filtered LFP, raster plots showing the spike patterns of 114 neurons, arrowheads indicating synchronous events in the raster plot, and animal running speed in a trial during the postlearning phase. Place cells were ordered by their place-field locations. (I) Spatial firing of a place cell with both stable and context-dependent properties. Spike positions are represented as red dots superimposed on the trajectories (gray). This cell showed stable spatial representations on path S-C_1_ and specifically fired when the rat approached C_1_ or C_2_. (J) Definition of context-dependent firing. The trajectories shown on the left and right are common as the rats move toward the check point (C_1_ or C_2_) and the goal, G, respectively. (K) Spatial firing rate distributions and the locations of place fields (shown as red *) of the place cell in I in the prelearning and learning phases are presented in the bottom and left panels, respectively. Joint place-field locations are plotted by black asterisks, representing the locations of all place-field pairs in different phases (green, stable; yellow, context-dependent). (L) All joint plots from all place cells. (M) Spatial correlation matrix of the population vector pairs at all location bins. (N) Summary of firing properties from all recorded cells (on, yellow; off, violet). (O) Bayesian decoding of rat trajectories from place cell spikes. Posterior probabilities of position estimates represented by a hot scale; position estimates overlaid with actual animal positions (cyan lines). (P) Errors of animal location estimates computed from datasets within each phase (cyan) or those averaged over all phases (magenta) by leave-one-out cross-validation: \**p* < 0.05, Mann-Whitney U test followed by Bonferroni correction. (Q) Average posterior probabilities of rat positions while running on a path as described above: the x-axis shows the phases of two theta cycles (white lines), and the y-axis shows the positions relative to the current animal’s location. (R) Comparison of quadrant scores of theta sequences across paths and phases: *\*p* < 0.05, Tukey test.

While the rats performed the task, the spike patterns of 355 neurons in the dorsal hippocampal CA1 region were recorded with independently movable tetrodes from the five rats (Fig. 1H and fig. S3). Some place cells had multiple place fields distributed throughout the environment (Fig. 1I-L and fig. S3E and 4F). Learning-induced changes in the place field locations were visualized by joint plots (Fig. 1K and 1 L and fig. S4A-C). The majority of place fields were stable irrespective of learning (green regions in Fig. 1K and 1 L); however, a subset of new fields emerged or disappeared in subsequent phases (fig. S4B). Notably, high density plots were detected for path S-C_1_ versus S-C_2_, path G-C_2_ versus S-C_2_, path S-C_1_ versus G-C_2_ (approaching check points), and path C_1_-G versus C_2_-G (approaching the goal area) (yellow regions). The same tendencies were confirmed by population vector correlation matrices (Fig. 1M and fig. S3B-D). The place fields that commonly emerged during an identical behavioral step, termed context-dependent fields (Fig. 1J), indicate the formation of generalized memory maps within the hippocampal circuit. The context-dependent codes observed here might consist partially of common mechanisms described by previous studies as goal-directed (*20*) or reward-predictive encoding (*21*). The proportions of place cells with stable and context-dependent place fields were 47.3% (112/237) and 48.9% (116/237), respectively (Fig. 1N and fig. S4E). Even when all these cell types were included, Bayesian decoding could still accurately reconstruct the rat’s actual positions from the position tuning curves of the cells (Fig. 1O and 1P; *p* < 0.05, Mann-Whitney U test versus shuffled data, U = 7317960132.5). Within a theta cycle, spatial firing of the place cells was organized in sequences (Fig. 1Q), the so-called theta sequences (*22*), which are considered a neuronal substrate that encodes immediate future and past locations and induces synaptic plasticity for memory acquisition during novel experiences (*23*). The quadrant scores, which represent the strength of the theta sequences, in path S-C_1_-G and the return path during the learning phase were significantly higher than those during the prelearning phase (Fig. 1R and fig. S5; *p* < 0.05, Tukey’s test). In the postlearning phase, the quadrant scores in path S-C_2_ remained significantly higher than those in the learning phase (*p* < 0.05, Tukey’s test). The stronger theta sequences possibly reflect increased cholinergic signals, which may lead to effective synaptic plasticity by coordinating with subsequent reward-related dopaminergic signals (*24*).

We next analyzed how new spatial learning is related to SWR-associated synchronous spikes of neuronal populations within a short (~100 ms) time window during stop periods, termed synchronous events (Fig. 2A and fig. S6). In the prelearning phase, synchronous events were temporally sparse, with frequencies below 0.4 Hz (Fig. 2B and fig. S7A-I). The frequencies of synchronous events and SWRs in the learning and postlearning phases became noticeably higher in the open field and goal area compared with the prelearning phase (Fig. 2B-D and fig. S7I; the distributions computed by the hierarchical Bayesian modeling with Markov chain Monte Carlo (MCMC) methods exhibited overlaps of less than 5%). On the next day (day 2) after learning, such SWR increases were no longer observed, and the frequencies were comparable to those in the prelearning phase (fig. S8), suggesting the necessity of novel learning to enrich synchronous events.

**Fig. 2.**
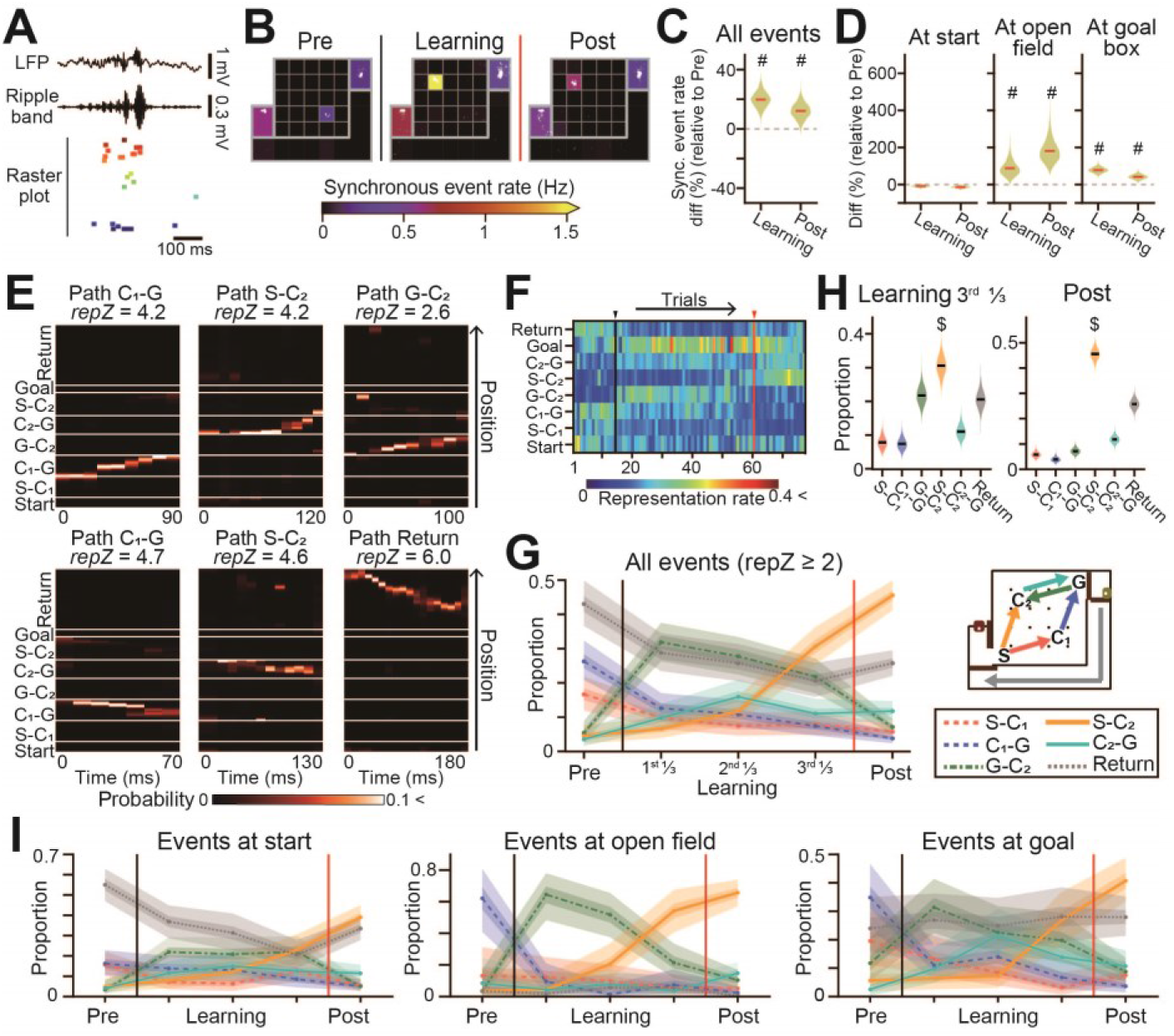
Prioritized representation of new episodes by learning-dependent synchronous events. (A) An SWR-associated synchronous event. (B) Pseudocolor maps of the frequencies of synchronous events. The synchronous event locations are indicated by superimposed white dots. (C) Distributions of the percentage of changes in synchronous events during the learning and postlearning phases compared to the prelearning phase. A pound sign (#) indicates that 0 is not in the 95% credible interval computed from the posterior probability distribution by MCMC. (D) Same as C, but separately plotted for the individual areas. (E) Bayesian decoding of animal trajectories from single synchronous events. Paths with the highest z-scored representation rates (*repZ*) are indicated above. (F) Color-coded matrices of representation rates for each path by rat. The black and red vertical lines indicate the reward replacement and learning points, respectively. (G) Learning-related changes in the proportions of represented paths by synchronous events. The learning phase was evenly divided into three phases (1^st^, 2^nd^, and 3^rd^ learning phases). The thick and thin shaded areas indicate the 50% and 95% credible intervals, respectively. (H) The data in G were specifically analyzed for the last 1/3 of the learning and postlearning phases. A dollar sign ($) indicates an overlap of less than 5% with any other distribution. (I) Same as G, but for separately plotted individual areas.

To examine the changes in neuronal ensemble patterns involved in synchronous events, correlation coefficients of vectorized population spikes were computed between synchronous events (fig. S9A). Event-to-event and trial-to-trial correlations changed substantially from learning and particularly increased around the learning points (fig. S9B-F), demonstrating that learning recruited new cell ensembles in synchronous events and that the stochasticity of such recruitment was reduced approximately when learning was established. To analyze the detailed episodes represented by synchronous events, Bayesian decoding was applied to estimate the animal trajectories (Fig. 2E and Supplementary Movie 1), and the representation rates (*reprates*) for the individual paths were computed as their posterior probabilities (Fig. 2F, fig. S10 and Supplementary Movie 2). For each synchronous event, a represented path was defined as the path with the highest *reprate*. In the prelearning phase, the majority of the represented paths were paths S-C_1_, C_1_-G, and the return path, corresponding with the actual animal trajectories. During the learning phase, paths G-C_2_ and C_2_-G became the majority of represented paths (Fig. 2G and fig. S10C), corresponding to the animal’s trial-and-error behavior along path G-C_2_-G (Fig. 1E-G). Notably, during the last 1/3 of the learning phase—before the rats finally settled on path S-C_2_-G—the largest fraction of synchronous events changed to represent path S-C_2_ (Fig. 2G and 2H; overlaps of the distributions computed by the hierarchical Bayesian modeling with MCMC methods were less than 5%), and such dynamic changes in representation patterns were more apparent in the open field than in the start and goal areas (Fig. 2I). These results demonstrate that future efficient behavioral episodes were already represented by synchronous events even before the animals changed their behavioral strategy. Another notable observation was that synchronous events more preferentially represented paths G-C_2_ and S-C_2_, where a reward (20 μl) newly emerged at the destination (C_2_), than path C_2_-G, where the a reward (200 μl) was continuously presented at the well-known destination (Goal area), which indicates that novel reward-related action policy, rather than the amount of reward, determines the priorities of episodes to be represented by hippocampal synchronous events (*14, 15*). Taken together, these results suggest that hippocampal synchronous events prioritize representations of salient episodes in reference to internal prediction errors and that such representation patterns undergo dynamic changes as learning proceeds.

To examine whether the learning-associated synchronous events corresponded with the replay events, the directionality and sequence strength of the spike trains for each synchronous event were assessed by computing a weighted correlation (*r*) and sequence score (*rZ*), respectively: synchronous events where |*r*| ≥ 0.5 were considered sequential events (Fig. 3A and fig. S12A-C). The participation rates of place cells with stable and context-dependent place fields in synchronous events were significantly higher than those of the other cells (Fig. 3B and fig. S11A-C; *p* < 0.05, Tukey’s test). In addition, these cell types contributed significantly to sequential events, as revealed by their significantly higher per-cell contributions (PCCs) (Fig. 3C and fig. S11D; *t*_*104*_ = 11.4, *t*_*107*_ = 9.8, *t*_*68*_ = 8.8, *t*_*145*_ = 9.1, *p* < 0.05, one-sample *t*-test versus 0). These results suggest that such stable and context-dependent representations may cooperatively contribute to creating learning-dependent synchronous events by utilizing a learned model for solving the task. In total, 345/2,586 (13.3%) and 390/2,586 (15.1%) synchronous events were classified as forward and reverse replay events, respectively (fig. S12A). Sequential events in the open field were biased toward forward rather than reverse directions in the prelearning phase, suggesting that forward representations primarily emerge in well-learned situations (Fig. 3D and fig. S12E; the overlaps between 0.5 and the distribution computed by the hierarchical Bayesian modeling with MCMC methods were less than 5%). Interestingly, this tendency was reversed after new learning was initiated; the reverse replays occupied significantly increased fractions in the latter learning phase, while the directionality of the replay events at the start and goal areas remained almost unchanged by learning. Overall, the sequential and trajectory events showed learning-dependent changes in the proportions of represented paths similar to those in Fig. 2G (Fig. 3E and 3F and fig. S12D and 13D), confirming that experience replays for the paths utilized in the new situation specifically developed with animal experience (Fig. 3G). Such learning-dependent changes in hippocampal replay patterns may be crucial for neuronal circuits to precisely reinforce new experiences that should be prioritized for learning.

**Fig. 3.**
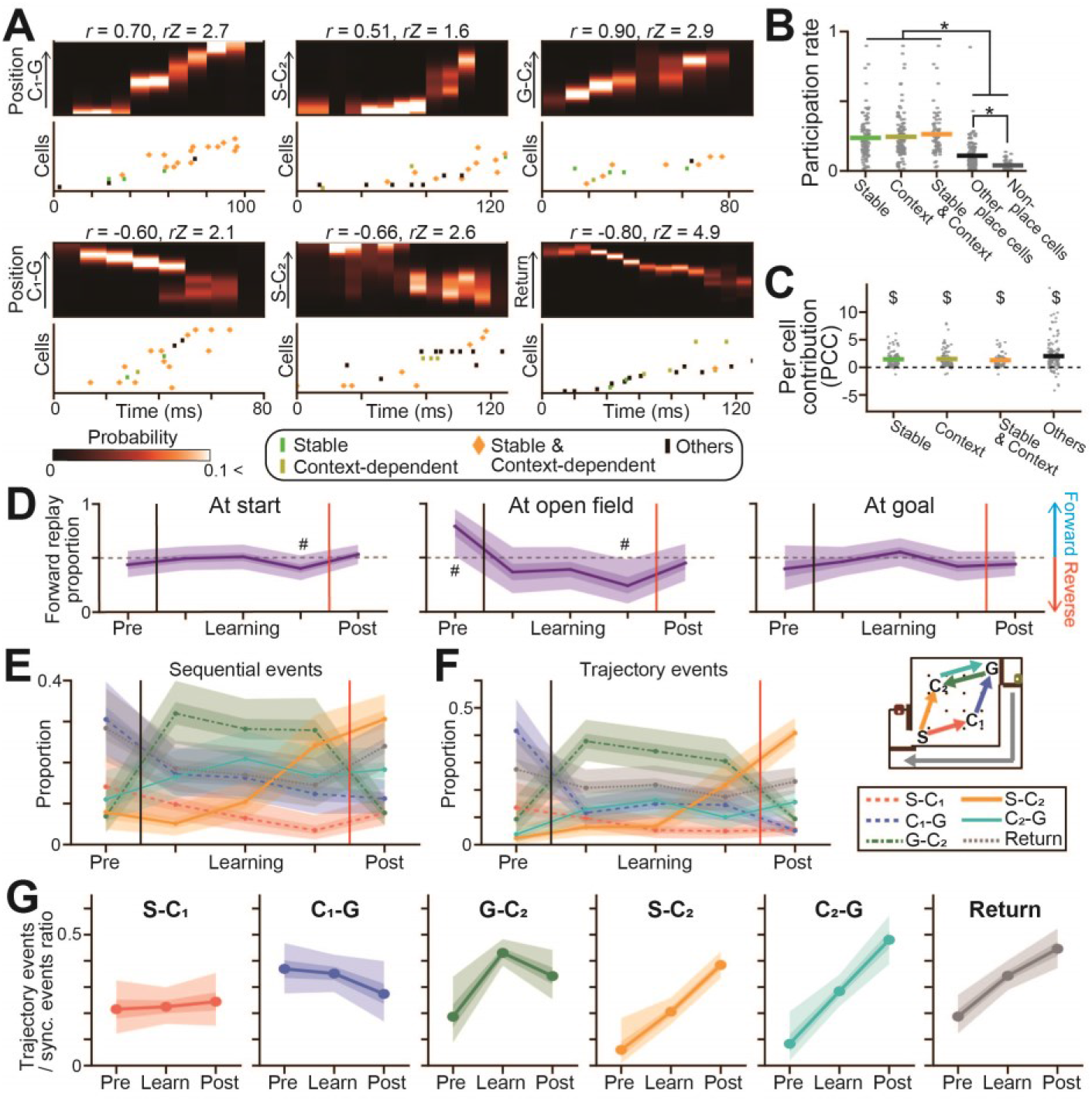
Representations by learning-dependent replay events. (A) Magnified views of the synchronous events shown in Fig. 2E and corresponding raster plots of the place cell spikes. The spikes are labeled based on cell type. (B) Participation rates of individual cells in synchronous events (*n* = 112, 116, 74, 154, and 47 cells). The gray dots represent each cell, and the colored lines represent the average: \**p* < 0.05, Tukey’s test. (C) Per-cell contribution of individual cells to replay events: $*p* < 0.05, one-sample *t*-test versus 0. (D) The directionalities of replay events in individual areas. The thick and thin shaded areas indicate the 50% and 95% credible intervals, respectively. A pound sign (#) indicates that 0.5 is not in the 95% credible interval. (E, F) Learning-related changes in the proportions of the represented paths by replay events with sequential and trajectory events. (G) The proportion of synchronous events assigned as trajectory events for each path. The thick- and thin_-shaded areas indicate the 50% and 95% credible intervals, respectively.

In order to test the causal role of SWR-associated synchronous events in learning performance, realtime disruption of SWRs was implemented by additionally implanting stimulation into the ventral hippocampal commissure (vHC); then, closed-loop feedback electrical stimulation with a single pulse (140-180 μA, 100 μs) was delivered upon the detection of hippocampal SWRs (Fig. 4A and 4B). As reported in earlier studies (*8, 9*), this manipulation can transiently eliminate SWR-associated neuronal events (Fig. 4C). Online disruption of ongoing SWRs after the replacement resulted in unstable animal trajectories across trials (fig. S14B and 14C), as demonstrated by a larger trajectory-to-trajectory distance (Fig. 4D and 4E) and significant increases in the probability of behavioral changes across two successive trials after the animals first took the most efficient path (i.e., start-S-C_2_) (Fig. 4F; *p* < 0.05, Tukey’s test). In contrast, stimulation delivered with a 250-ms delay relative to SWRs (delayed control) and feedback stimulation in a nonlearning control group did not evoke such behavioral changes (Fig. 4E and 4F; *p* > 0.05, Tukey’s test). The SWR disruption-induced behavior deficit implies that learning-dependent SWRs and replay events are necessary to support efficient learning and reinforce specific learned behavior.

**Fig. 4.**
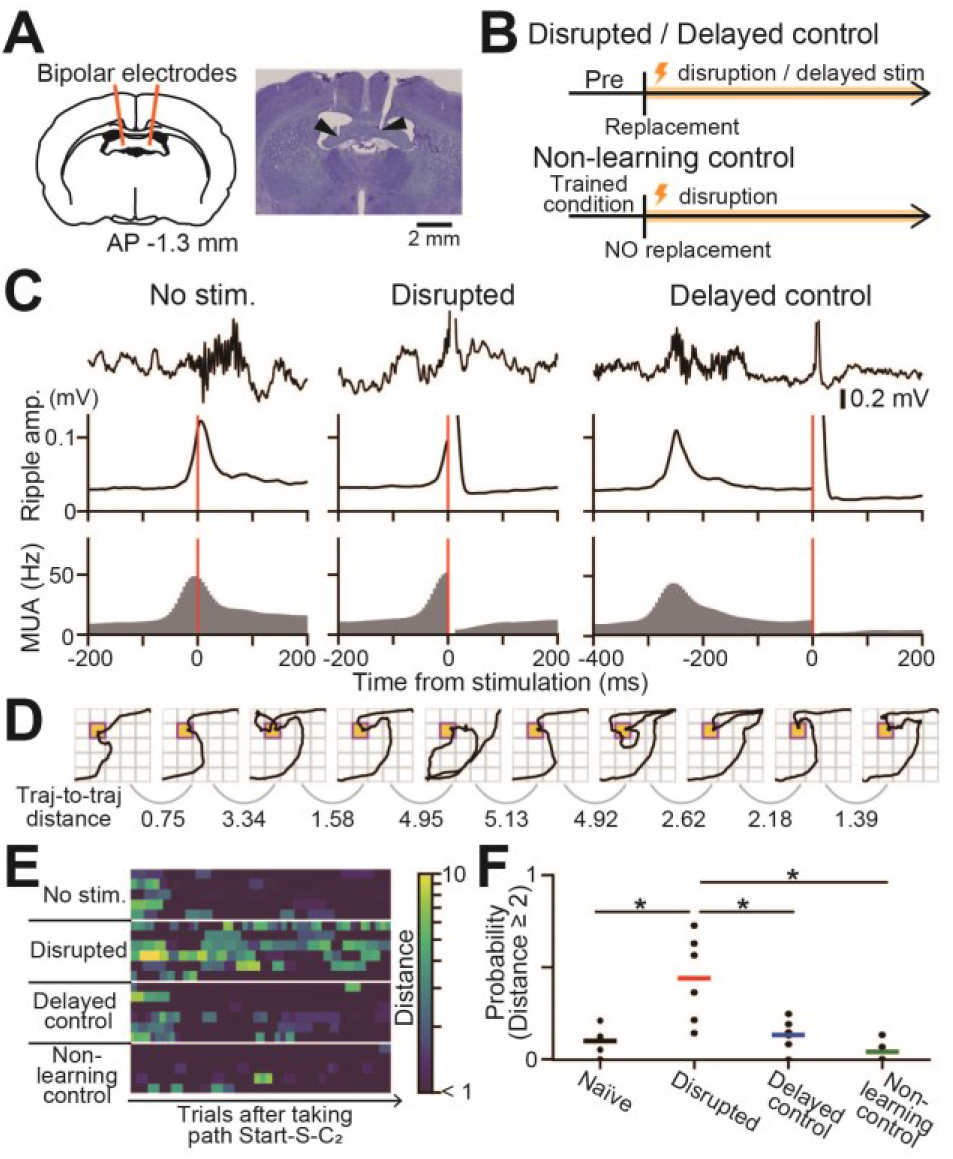
Impairment of learning by inhibition of SWRs. (A) Electrodes were bilaterally implanted in the vHC, and real-time closed-loop electrical stimulation was applied to disrupt hippocampal SWRs. (B) Schematic of the experiments. Timed or 250-ms delayed stimulation was applied upon SWR detection after reward relocation (disrupted or delayed control). In the nonlearning control group, the reward point was not relocated, but feedback stimulation was applied. (C) Original LFP traces, ripple power, and multiunit firing rates aligned to the time of stimulation in the unstimulated, disrupted, and delayed control groups. The red lines denote the time of stimulation. (D) Representative trajectories in a rat with disrupted SWRs. Trajectory-to-trajectory distances were computed between all pairs of successive trials. (E) Pseudocolor maps of the trajectory-to-trajectory distance for all individual rats after the animals first took the path start-S-C_2_. (F) The probability of trials with an observed distance ≥ 2 after the animals first took the path start-S-C_2_: *\*p* < 0.05, Tukey test.

Our finding of context-dependent encoding by certain hippocampal cell ensembles is consistent with the theory of a predictive map (*2*) and suggests that hippocampal neurons may even represent generalized behavioral procedures (e.g., moving toward a checkpoint) and process abstract factorized codes (*19*). The potential ideas could extend a predictive map that has been primarily considered a tool for spatial information processing (*2, 16*) to a predictive model that can process more abstract and contextual information, which could further enrich predictive computations of task state values and support model-based behavior based on past episodes (*25*). Moreover, we demonstrated that, as the animals learned and developed new trajectory patterns in response to reward replacement, hippocampal cell ensembles prominently increased theta sequences and SWR-associated replay events, both of which have been considered neurophysiological mechanisms to induce circuit plasticity (*3, 9, 23, 24*). These observations suggest that the hippocampal predictive map can be flexibly reorganized to properly represent varying task states when animals learn new behavioral policies in reference to their past episodes (*2, 16, 26, 27*). Moreover, we demonstrated that increased hippocampal replay events after reward replacement preferentially represented the most salient (C_2_-related) experiences and strategy that needed to be learned and exploited by the animals. Such prioritized replays of salient experiences may be effective to amplify and consolidate neuronal ensembles that encoded newly learned experiences onto the hippocampal circuit. Concordantly, recent studies of artificial intelligence have demonstrated that incorporating “prioritized experience replays” and episodic meta-learning into deep neural networks could improve the efficiency of reinforcement learning in artificial agents (*11, 14, 17, 26, 28*). Taken together, the evidence suggests that forming an effective predictive map and rehearsing salient external interactions by prioritized replays in the internal neuronal network are common beneficial mechanisms for both living brains and artificial agents to learn and reinforce specific behavioral strategies.

## Author contributions

H.I. and T.S. designed the study. H.I. performed the surgery, acquired the electrophysiological data, and performed the analysis. Y.I. supervised the project. H.I. prepared all figures. H.I. and T.S. wrote the main manuscript text, and all authors reviewed the main manuscript text.

## Data and materials availability

All data are available in the main text or the supplementary materials.

## Supplementary Materials

### Materials and Methods

#### Approvals

All the experiments were performed with the approval of the experimental animal ethics committee at the University of Tokyo (approval number: P29-7) and conducted according to the NIH guidelines for the care and use of animals.

#### Subjects

A total of 17 male Long Evans rats (3-6 months old) with preoperative weights of 400-500 g were used in this study. The animals were housed individually and maintained on a 12-h light/12-h dark schedule with lights off at 7:00 AM. All the animal subjects were purchased from SLC (Shizuoka, Japan). Following at least 1 week of laboratory adaptation, the rats were reduced to 85% of their *ad libitum* weight by limiting daily feeding. Water was readily available.

#### Behavioral apparatus for a spatial learning task

The overall size of the platform used for the spatial learning task was 1.2 m × 1.4 m, elevated 70 cm from the floor, with a wall height of 30 cm. This platform contained a square (1 m per side) open field, a 20 cm × 40 cm start area, a 20 cm × 30 cm goal area, and an L-shaped outer peripheral alleyway connecting the start and goal areas (return path) with a width of 20 cm (Fig. 1A and Supplementary Fig. 1B). The open field was evenly divided into 5 × 5 grids, and a black pole with a height of 10 cm and a diameter of 2 cm was placed at each grid point. The borders between the start area and the open field, the open field and the goal area, the goal area and the peripheral alleyway, and the peripheral alleyway and the goal were partitioned by automatic doors, termed door 1, door 2, door 3, and door 4, respectively (Fig. 1A and Supplementary Fig. 1B). All the doors were semiautomatically controlled. The other areas were partitioned by walls with a height of 22.5 cm. The floor, doors, and walls were all made of ABS resin. In the start area, a nose-poke reward port (3 cm in diameter) was attached to one side of the wall, and a white LED (5 × 5 mm) was attached 12 cm above the port. A speaker for sound presentation was placed outside the platform. In the goal area, an automatic reward-feeding port (7 × 6 cm) was attached to one side of the wall.

#### Trainings before surgery

All the behavioral experiments occurred in the dark phase with a light intensity of 1 lux. Training consisted of several steps (Supplementary Fig. 1C). On the first 2 days, each rat was habituated to the open field by allowing it to freely forage for randomly scattered chocolate milk for 10 min (Supplementary Fig. 1C, termed habituation).

After the habituation period, each rat was trained to perform voluntary nose poking in the start area for 2 days (Supplementary Fig. 1C, termed “nose-poke training”). Continuous nose poking for 1 s triggered 5-kHz sounds at 10 Hz for 0.3 s and a subsequent 10-kHz sound for 0.7 s. During the sound presentation period, the rat could obtain 25 μl of 30% sucrose eight times dispensed from the port by a syringe pump. Lighting of the white LED indicated that a reward was available at the port, and the LED was turned off when the rat could perform valid nose poking. After reward consumption, the rat had to leave the port for at least 2 s to initiate the next nose poking. When the intertrial intervals terminated after the rat left the port, the LED was again turned on to initiate the next trial.

Next, each rat was trained to run in the field (Supplementary Fig. 1C, termed “running training”). Similar to the nose-poke training, a rat initiated a trial by nose poking in the start area and obtained sucrose water during cue-sound presentation. Ten seconds after the onset of the sound presentation, the door between the start area and open field (door 1) was automatically opened, allowing the rat to enter the field. At the same time, 20 μl of chocolate milk reward was placed in the first lattice after the start area (S). When the rat entered lattice S, 5-kHz cue sounds at 10 Hz for 0.3 s followed by continuous 10-kHz cue sounds were presented until the rat reached the goal area. When the rat reached the last lattice before the goal area (lattice G), the door between the field and goal area (door 2) was opened so that the rat could enter the goal area. In the goal area, the rat received 200 μl of chocolate milk reward dispensed from the port by a syringe pump. Twenty seconds after the onset of entry into the goal area, the doors between the goal area and the peripheral alleyway (door 3) and between the peripheral alleyway and the start area (door 4) were opened, allowing the rat return to the start area through the alleyway to complete the trial (Supplementary Fig. 1D). During this running training, rats could follow any trajectory through the field from the start area to the goal area. On average, it took well-trained rats approximately 1 min to complete one trial. If a rat stayed in the field for more than 2 min without opening door 2, door 2 was opened, irrespective of the rat’s trajectory, so that the rat could enter the goal area. This training was repeated daily until the rat was able to complete more than 7 trials from the start to the goal area within 10 min. To achieve this performance criterion required between 5 and 7 days. After each rat met this criterion, it underwent surgery for electrode implantation.

To detect nose poking and reward consumption, infrared photoreflectors ware attached to the start and goal areas. To monitor the rat’s moment-to-moment position, three red LEDs with a diameter of 5 mm were attached to the rat’s back with a harness, and the positions of the LED signal were automatically tracked in real time at 24.5 Hz using a video camera attached to the ceiling. Depending on the instantaneous rat positions and sensor inputs, door openings and reward feedings were automatically regulated; these were digitized and timestamped by a laptop computer. Because all the apparatuses were controlled by a computer, it was not necessary for an experimenter to handle the rat after a task began.

#### Training for a spatial learning task after surgery

After recovery from surgery, the rats again underwent running training for 1-2 days (Supplementary Fig. 1C). After confirming that the rats had again reached the criterion for performance, as in the presurgery period, the rats were trained in the C_1_ condition for 2-3 days (Supplementary Fig. 1C). Similar to the running training, a rat initiated a trial by nose poking and obtained sucrose water during cue-sound presentation. Ten seconds after the onset of the sound presentation, the door between the start area and open field (door 1) was opened automatically, allowing the rat to enter the field. At the same time, 20 μl of chocolate milk reward was placed in the lattice fourth from left and second from bottom, termed check point 1 (C_1_). The rat was trained to run from the first lattice after the start area (S) to C_1_ (path S-C_1_), obtain the reward at C_1_ and then run from C_1_ to the last lattice before the goal area (G) (path C_1_-G). When the rat entered C_1_, 5-kHz cue sounds were presented at 10 Hz for 0.3 s followed by continuous 10-kHz cue sounds until the rat reached the goal area. This sound cue helped a rat recognize that its current state was correct for obtaining chocolate milk in the goal area. When the rat reached G after passing through C_1_, the door between the field and the goal area (door 2) was opened so that the rat could enter the goal area. In the goal area, the rat obtained 200 μl of chocolate milk reward. Twenty seconds after the onset of reward dispensation, the doors between the goal area and peripheral alleyway (door 3) and the between the peripheral alleyway and start area (door 4) were opened, allowing the rat to return to the start area through the alleyway to complete the trial. The next trial started when the rat again poked the reward port in the start area. At that point, all the doors were closed to initiate the next trial. In some cases, this training was performed with a recording headset and cable attached so that the animals became familiar with the recording condition.

To monitor the rat’s moment-to-moment position, three red LEDs were attached to the electrode assembly, and the position of the LED signal was automatically tracked in real time at 24.5 Hz using a video camera attached to the ceiling. On all training and recording days, the rats were kept in a rest box (33 × 33 cm) outside the field for tens of minutes before and after performing the task.

#### A spatial learning task on a recording day

On a recording day, the rats first performed the same task with a reward placed on C_1_, termed the prelearning phase. After several trials, the rewarded check point was moved from C_1_ to the second lattice from the left and fourth from the bottom, termed check point 2 (C_2_). In this phase, when the rat visited C_2_, 5-kHz cue were played at 10 Hz for 0.3 s followed by continuous 10-kHz cue sounds were presented until the rat reached the goal area. The chocolate milk reward volume and all the other task conditions were similar to those in the prelearning phase. In this situation, the most efficient behavioral strategy was to run directly from S to C_2_ (path S-C_2_), take the reward placed on C_2_, and then run from C_2_ to G (path C_2_-G). After the reward replacement, the rats first exhibited trial-and-error behavior for several attempts to find an efficient trajectory, but they gradually learned to take the most efficient trajectory: path S-C_2_-G. The detailed definition of a learning point is provided below.

#### Surgical procedures

Five and twelve rats underwent surgery to implant recording electrodes only and a combination of recording and stimulating electrodes, respectively. Briefly, the rats were anesthetized with isoflurane gas (0.5-2.5%), and a 2-cm midline incision was made from the area between the eyes to the cerebellum. For 5 rats, a craniotomy with a diameter of 0.9-1.6 mm was created above the right dorsal hippocampus (3.8 mm posterior and 2.8 mm lateral to bregma) using a high-speed drill, and the dura was surgically removed. Two stainless-steel screws were implanted in the bone above the prefrontal cortex to serve as ground electrodes. Using a 3D printer (Form 2, Formlabs) an electrode assembly consisting of 16 independently movable tetrodes was created and stereotaxically implanted above the craniotomy. The tips of the tetrode bundles were lowered to the cortical surface, and the electrodes were inserted 1.0 mm into the brain at the end of the surgery. The electrodes were constructed from 17-μm-wide polyimide-coated platinum-iridium (90/10%) wire (California Fine Wire California Fine Wire Co., Grover Beach, CA), and the electrode tips were plated with platinum to reduce their electrode impedances to 150-300 kΩ at 1 kHz. For 12 rats, an electrode assembly that consisted of 7 independently movable tetrodes was implanted using the same procedures as described above. In addition, the craniotomies (1.3 mm posterior and 1.7 mm lateral to the bregma) with a diameter of ~1 mm were created using a high-speed drill, and stainless bipolar electrodes were implanted at a depth of 3.7 mm at an angle of 6.9° into the right side or both sides of the ventral hippocampal commissure (vHC).

All the recording devices were secured to the skull using stainless-steel screws and dental cement (Provinice, Shofu Inc., Kyoto, Japan). Following surgery, each rat was housed individually in transparent Plexiglass with free access to water and food for at least 5 days and was then food-deprived until they reached 85% of their previous body weight.

#### Adjusting electrode depth

Each rat was connected to the recording equipment via a Cereplex M (Blackrock) digitally programmable amplifier, close to the rat’s head. The output of the headstage was connected via a lightweight multiwire tether and a commutator to a Cerebus recording system (Blackrock), a data acquisition system. Electrode turning was performed while the rat was resting in a pot placed on a pedestal. The electrode tips were slowly advanced by 25-100 μm per day for 11-24 days until spiking cells were encountered in the CA1 layer of the hippocampus, which was identified on the basis of local field potential (LFP) signals and single-unit spike patterns. After the tetrodes were adjacent to the cell layer, as indicated by the presence of multiunit activity, the tetrodes were settled into the cell layer for stable recordings. Subsequently, recordings were conducted over a period of several days as described below.

#### Electrophysiological recording

Electrophysiological data were sampled at 2 kHz and low-pass filtered at 500 Hz. Unit activity was amplified and high-pass filtered at 750 Hz. Spike waveforms above a trigger threshold (−50 μV) were timestamped and recorded at 30 kHz for 1.6 ms.

#### Closed-loop electrical stimulation

Upon the online detection of SWRs, closed-loop electrical stimulation was performed using extension code implemented on the Cerebus recording system (Blackrock) and custom-created C code. A tetrode implanted into the hippocampus was chosen, and the envelope of its bandpass (100-400 Hz)-filtered LFP signals at 30 kHz was estimated in real time as described previously (*8*). The smoothed estimate of the envelope (*env*_*est*_) of filtered LFP signals was computed as follows:

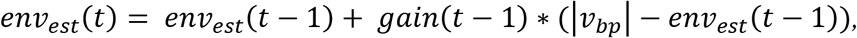

where

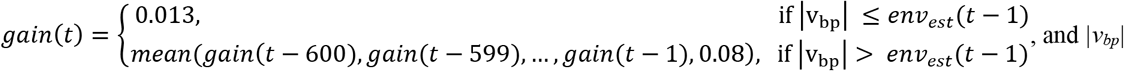

denotes the absolute values of the filtered LFP signals. The length of the gain buffer was set to 600 (20 ms). Estimated values of smoothed mean (*mean*_*est*_) and standard deviation (*std*_*est*_) were then computed as follows:

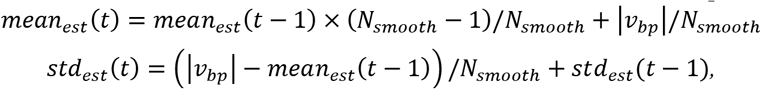

where *N*_*smooth*_ was the number of samples for smoothing (typically, set to be 150,000 in 500 ms). SWRs were detected online when the animal’s running speed was below 5 cm/s and when the envelope exceeded the detection threshold of 3-4 standard deviations above the estimated mean computed from LFP signals during periods in the rest box. At the time of SWR detection, an electrical pulse with a duration of 100 μs and an amplitude of 140-180 μA was applied to the vHC; the stimulation rate was limited to a maximum of 4 Hz. For delayed control stimulation, stimulation was applied with a latency of 250 ms after the onset of ripple detection so that the stimulation occurred outside the detected SWRs.

#### Histological analysis to confirm tetrode locations

After the experiments, the rats received an overdose of urethane and were intracardially perfused with 4% paraformaldehyde in PBS and decapitated. To aid in the reconstruction of the electrode tracks, the electrodes were not withdrawn from the brains until more than 3-4 hours after perfusion. After dissection, the brains were fixed overnight in 4% paraformaldehyde (PFA) and then equilibrated with a sequence of 20% sucrose and 30% sucrose in PBS. Frozen coronal slices (50 μm) were cut using a microtome (Sliding Microtome, SM2010 R, Leica Biosystems, Wetzlar, Germany), and serial sections were mounted and processed for cresyl violet staining. To perform cresyl violet staining, the slices were rinsed in water, counterstained with cresyl violet, and coverslipped with hydrophobic mounting medium (PARAmount-D, Falma, Tokyo, Japan). The positions of all the tetrodes were confirmed by identifying the corresponding electrode tracks in histological tissue with an optical microscope (All-in-One Fluorescence Microscope BZ-X710, Keyence Corporation, Osaka, Japan).

#### Spike sorting of hippocampal neurons

Spike sorting was performed offline using the graphical cluster-cutting software MClust (*29*). Rest recordings before and after the behavioral paradigms were included in the analysis to assure recording stability throughout the experiment and to identify hippocampal cells that were silent during the running behavior. Clustering was performed manually in two-dimensional projections of the multidimensional parameter space (i.e., comparisons between waveform amplitudes, the peak-to-trough amplitude differences, waveform energies, and the principal components of waveforms, each measured on the four channels of each tetrode). Autocorrelation and cross-correlation functions were used as additional separation criteria. Refractory periods of spikes were considered to increase confidence in the successful isolation of cells. Clustering quality was measured by computing the L_ratio_ and isolation distance (*30*). A cluster was considered to be a cell when the L_ratio_ was less than 0.40. Overall, 114, 72, 68, 79, and 22 cells were recorded from rat 1, 2, 3, 4, and 5, respectively.

#### Analysis of animal trajectories and definition of a learning point

To analyze animal trajectory patterns, the open field was evenly divided into a 5 × 5 lattice and each grid cell was labeled with character strings “a-y” (Supplementary Fig. 1E). Five paths connecting specific pairs of four lattices (u (S), e (G), s (C_1_), and g (C_2_)), S-C_1_, C_1_-G, G-C_2_, C_2_-G, and S-C_2_, were analyzed. The minimum string length of all these paths was 5. An animal’s trajectory was divided into trajectory segments connecting two of the four specific lattices, and each trajectory segment was classified into one of the five paths if the length of the trajectory segment was less than 8 (an example shown in Supplementary Fig. 1G; red, S-C_1_; blue, C_1_-G; green G-C_2_; cyan, C_2_-G; orange, S-C_2_). Trajectory segments that did not meet this criterion were classified as “other”. The majority of trajectory segments were classified into paths S-C_1_, C_1_-G, G-C_2_, S-C_2_, and C_2_-G (for more detail, see Fig. 1E and 1F). In all the following analyses, these five paths were selectively analyzed unless otherwise specified. For the process of linearizing the rat trajectories on individual paths, see Supplementary Fig. 1I.

For each rat, a learning point (between the learning and postlearning phases) was computed by depicting a learning curve with a moving average window of 5 trials (Supplementary Fig. 2B and 2D). In the learning phase, the optimized trajectory was defined as path S-C_2_-G with a total string length less than 12. In a learning curve, a learning point was defined as one trial before the point where the optimized trajectory percentage first exceeded 50%. If that trial did not include the optimized trajectory, the learning point was defined as the next trial.

#### Place field characteristics

For each cell, an average spatial firing-rate distribution at an animal’s running speed of more than 5 cm/s was computed along the linearized trajectories (Supplementary Fig. 1I) with a bin size of 2 cm; these data were then smoothed by a Gaussian filter (σ = 5 cm). For each distribution, a null distribution was constructed using an identical procedure, in which the spike locations were randomized while the total number of spikes was preserved. For one original distribution, 1,000 null distributions were created. The null distributions were created separately for the open field and the return path. The candidate position bins for the place fields were detected where the original firing rates exceeded the 99th percentile of those in the 1,000 null distributions. Place fields were defined as candidate position bins when more than five candidate position bins with firing rates greater than 1 Hz were detected consecutively as described previously (*31*). For each place field, the place field center was defined as the position with the maximum spiking rate. The place field detection was performed separately for each area and path (start area, S-C_1_, C_1_-G, G-C_2_, S-C_2_, C_2_-G, goal area, and return). Neurons with at least one place field in the open field or return path in at least one of the three phases were classified as “place cells”.

The characteristics of each place cell’s place fields on the five paths (S-C_1_, C_1_-G, G-C_2_, S-C_2_, and C_2_-G) were defined by depicting joint plots of place field locations between all phase pairs: prelearning versus learning phases, learning versus postlearning phases, and prelearning versus postlearning phases (an example cell is shown in Fig. 1K, 1 L and all cells are shown in Supplementary Fig. 4). Place fields that emerged or disappeared in a later phase were classified as “appearing” or “disappearing”, respectively. A place field that emerged at the same location with an interfield interval of less than 20 cm across two phases was classified as “stable”. Place fields that commonly emerged when an animal approached or left specific lattices (S, C_1_/C_2_, and G) were classified as “context-dependent”. A place field that emerged at the same location irrespective of the running direction on paths C_2_-G and G-C_2_ was classified as “bidirectional”. Place cells with these place field characteristics are summarized in Fig. 1N and Supplementary Fig. 4D.

#### Spatial correlation of place-cell population vectors

In each place cell, a spatial firing-rate distribution in each phase was normalized so that the maximum firing rate was 1, as shown in Supplementary Fig. 3A. In each phase, a population vector with a location bin size of approximately 2 cm was constructed from the normalized firing-rate distributions of all the recorded place cells. Correlation coefficients for all pairs of population vectors were computed to construct a population vector correlation matrix (Supplementary Fig. 3B and 3C) as follows:

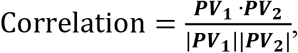

where ***PV*** is a population vector in each phase.

#### Bayesian decoding of animal locations

To reduce memory requirements, uniform prior Bayesian decoding was applied to estimate the animals’ positions from the spike trains. The spatial firing-rate distributions of individual place cells were used as position-tuning curves. As described previously (*5*), assuming Poisson firing statistics and a uniform prior over position, the posterior probability of the animal’s location (*loc*) in a time window (*τ*) including neuronal spike patterns (*s*) was computed as follows:

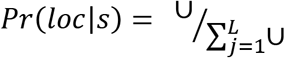

where

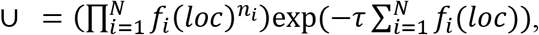

*f*_*i*_(*loc*) is the position tuning curve of the i-th neuron, and *N* and *L* are the total numbers of neurons and total location bins, respectively. The time window *τ* was set to 490 ms (12 video frames), 20 ms, and 21 ms (1/6 theta cycles) to estimate the animal’s positions at timescales of behavior (Fig. 1), synchronous events (Fig. 2E and 3A), and theta cycles (Fig. 1Q), respectively. For visualization purposes and trajectory event detection, decoding for synchronous events was conducted with a 10-ms sliding window (Fig. 2E).

Fig. 1P shows a decoding accuracy comparison between the original position tuning curves and those from the surrogate position tuning curves in which the spatial selectivity of place cells was randomized. To create the surrogate position tuning curves, individual firing rates in the spatial firing-rate distributions were randomly shifted across all location bins for each cell before applying the Gaussian filter; then, the shuffled spatial firing-rate distributions were smoothed with a Gaussian filter (σ = 5 cm).

#### Theta sequence

LFP signals at a running speed of more than 5 cm/s were bandpass-filtered at the theta (6-10 Hz) band, and the peak and trough in each theta cycle were detected. Each theta cycle was evenly divided into 6 bins with a time window of ~21 ms, and Bayesian decoding of the animal’s positions relative to the current locations was performed in this time window as described above. The data obtained when the animals were located within 15 cm of the start and end points in each path were excluded from this analysis. The decoded probabilities (80 cm around the animal’s location and 2⁄3 of the theta cycle around the center of theta cycles) were divided into four quadrants, and a quadrant score was computed as the difference in the average decoded probabilities between the second and fourth quadrants and the first and third quadrants (Fig. 1R), as described previously (*32*).

#### Detection of synchronous event

For each rat, the instantaneous spike rates (bin = 1 ms) were averaged over all the recorded neurons and smoothed with a Gaussian filter (σ = 15 ms). In addition, the mean and standard deviation (SD) of instantaneous spike rates during stop periods with a running speed of less than 5 cm/s were computed from all the recorded neurons, termed the *mean*_*base rate*_ and *SD*_*base rate*_, respectively (Supplementary Fig. 9C, left). Using these baseline variables, the instantaneous spike rates were z-scored as shown in the bottom trace in Supplementary Fig. 9A and Supplementary Fig. 9C. Candidate synchronous events were detected when the z-scored spike rates exceeded 2 and the number of active neurons exceeded 4. The onset and offset of candidate synchronous events were marked at the time points when the z-scored spike rates first exceeded and fell below 0, respectively. Candidate synchronous events in which the duration was less than 50 ms or more than 2000 ms were excluded from further analyses.

Next, the overall ripple power during candidate synchronous events was computed. In each tetrode, the LFP signals were bandpass filtered at 150-250 Hz, and the envelope of the filtered LFP traces was computed via the Hilbert transformation. Then, the envelope was smoothed with a Gaussian filter (σ = 4 ms). In addition, baseline mean and SD vectors of the smoothed envelopes during the stop periods were computed in each tetrode, termed ***mean***_***base power***_ and ***SD***_***base power***_, respectively. Based on these variables, the Mahalanobis distance was computed at each time point, termed overall ripple power.

Using the Gaussian mixture model (GMM), the distributions of the z-scored spike rate and z-scored overall ripple power in all candidate synchronous events were each divided into two distributions, as shown in Supplementary Fig. 9E. Synchronous events were detected when time points were included in the larger distributions of z-scored spike rate or z-scored overall ripple power (colored in magenta in Supplementary Fig. 9E).

#### Detection of SWRs

In a tetrode, LFP signals were bandpass filtered at 150-250 Hz, and the envelope of the filtered LFP traces was computed via the Hilbert transformation. The envelope was then smoothed with a Gaussian filter (σ = 4 ms), termed ripple power. In addition, the mean and SD of ripple power during stop periods with a running speed of less than 5 cm in the task periods were computed in each tetrode, termed *mean*_*base power*_ and *SD*_*base power*_, respectively. SWR events were detected when the envelope exceeded (*mean*_*base power*_ + 3 × *SD*_*base power*_). The onset and offset of SWR events were marked at the time points when the ripple power first exceeded and fell below *mean*_*base power*_, respectively. SWR events with a duration less than 50 ms or more than 500 ms were excluded.

#### Similarity of synchronous events

To quantify the similarity of the synchronous events, each synchronous event was converted to an *N*-dimensional vector containing entries of the spike counts of individual neurons during the event, where *N* denotes the total number of neurons. Correlation coefficients between all possible vector pairs were calculated to construct an event-to-event correlation matrix (Supplementary Fig. 9A). A trial-to-trial correlation matrix was then constructed from the event-to-event matrix by calculating the average of all the correlation coefficients included in each trial pair except for the values on the diagonal line. Trials with fewer than 2 synchronous events were not analyzed. To evaluate the significance of the correlation coefficients in the trial-to-trial correlation matrix, the same analysis was applied to randomized data, in which the temporal order of the actual synchronous events in the original data was shuffled while preserving the total numbers of synchronous events in the individual trials. This randomization process was repeated 1,000 times. At each point, a z-scored correlation coefficient of the real data was computed from the distribution of the 1,000 surrogate correlation coefficients (Supplementary Fig. 9B, bottom). A z-scored phase-to-phase correlation matrix was computed, similar to trial-to-trial correlations (Supplementary Fig. 9E and 9F).

#### Changes in the frequency of synchronous events and SWRs

Similar to the work of (*15*), hierarchical Bayesian modeling was applied to estimate the posterior probability of the number of synchronous events with the parameters estimated by Markov chain Monte Carlo (MCMC) methods. We assumed that synchronous events emerge according to Poisson firing statistics and varied across individual animals, as follows:

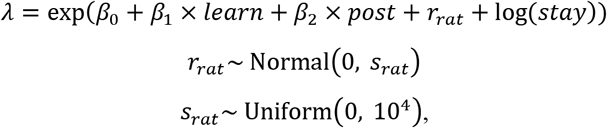

where *λ* is the expected value of the number of synchronous events, *stay* is the duration (s) with a running speed of less than 5 cm, *learn* and *post* are dummy variables for the learning and postlearning phases, respectively. Here, *r*_*rat*_ reflects individual rat differences. Based on the estimated coefficients, the percentage of changes in the frequency of synchronous events for each trial in the learning and postlearning phases relative to those in the prelearning phase were computed as 100 × [exp(β_*i*_) − 1], where *i* = 1 or 2, representing the learning or postlearning phases, respectively (Fig. 2C and 2D). Similarly, in Supplementary Fig. 7C, the percentage of changes in the number of synchronous events in each trial was computed with log(*stay*) = 0.

Similar to the synchronous events, the changes in SWR frequencies were quantified in Supplementary Fig. 7G, 7I, 8E, and 8G.

#### Representation rate

For each synchronous event, the posterior probability of the animal’s location was computed by Bayesian decoding as described above (examples shown in Fig. 2E). The posterior probabilities within individual paths were averaged over the all location bins and summed over all the time bins. The summed posterior probabilities for the individual paths were then normalized so that the sum of the probabilities over all paths was 1, termed “representation rates”. To depict the color-coded matrix in Fig. 2F, the representation rates of individual paths were averaged over each trial. For each synchronous event, a z-scored representation rate (*repZ*) was computed for the path segment with the highest representation rate as follows:

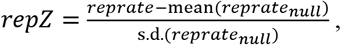

where *reprate* denotes a representation rate for the path segment and *reprate*_*null*_ denotes the representation rates obtained from the decoding of the 1,000 surrogate synchronous events where the randomized position-tuning curves were created using the procedure described above.

#### Synchronous events representing individual path segments

Specifically, paths S-C_1_, C_1_-G, G-C_2_, C_2_-G, S-C_2_, and the return path were analyzed. For each synchronous event, the representative paths were defined as the paths with the highest *reprate*. Hierarchical Bayesian modeling was applied to estimate the posterior probability of the proportions of represented paths with the parameters estimated by Markov chain Monte Carlo (MCMC) methods. We assumed that replays for the individual paths in each trial emerged according to a multinomial distribution as follows:

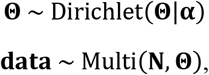

where **data** was a *T* × *U*-dimensional vector matrix whose entries were the number of represented paths in each trial, *T* and *U* were the number of trials and paths (*U* = 6), respectively, ***N*** was a *T* × 1-dimensional vector containing the total number of represented paths in each trial, and the parameters **Θ** and **α** were *U* × 1-dimensional vectors for multinomial distribution and Dirichlet distribution, respectively. The initial vector **α** was set to [1,1,1,1,1,1,1]. The parameter **Θ** for the events where *repZ* ≥ 2 was plotted as the proportions of represented paths in Fig. 2G, 2H, and 2I. The proportions of represented paths for all events are shown in Supplementary Fig. 10C and 10D.

#### Sequence scores of sequential events

Sequence scores were defined for individual synchronous events as described previously (*31*). For sequential events and trajectory events, the data from rat 1, 2, 3, and 4 were analyzed (with 68-114 cells). The path with the highest representation rate was analyzed. Replays for the start and goal areas were excluded from the analysis. For the represented path segments, *Pr*(*loc*|*s*) constructed from a synchronous event was smoothed over time bins with a Gaussian filter (σ = 10 ms). In the filtered *Pr*(*loc*|*s*), a maximum a posteriori probability (MAP) was computed as the largest posterior probability across all positions per time bin(*33*), and the time bins with MAPs greater than (5 × 1/*n*_*loc bin*_) were included for the analysis, where *n*_*time bin*_ and *n*_*loc bin*_ were the numbers of time and location bins in the synchronous event, respectively. If the number of time bins in which MAP exceeded the threshold was less than 3, all the time bins were analyzed. *Pr*(*loc*|*s*) was normalized within the represented paths as follows:

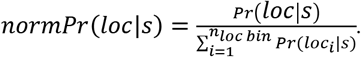

A sequence score *r(loc, time; normPr)* representing the weighted correlation between time and location was computed as follows:

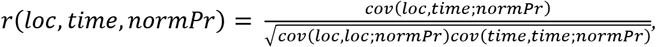

where

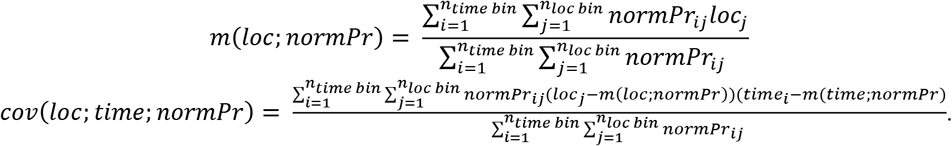

For each sequence score *r*, a z-scored sequence score (*rZ*) was computed as follows:

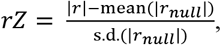

where *r*_*null*_ is the sequence score obtained from 1,000 surrogate synchronous events with randomized position tuning curves, which were randomly shifted within the represented path for each cell. Distributions of *r* and *rZ* values are presented in Supplementary Fig. 12A. Synchronous events with an |*r*| ≥ 0.5 were considered sequential events, and the replay directions were determined by the signs of the correlations *r*, where positive and negative correlations of *r* represent forward and backward replay directions, respectively.

#### Per cell contribution (PCC)

To quantify the contribution of each cell type (stable, context-dependent, and other place cells) to the sequence scores of synchronous events, PCCs were computed as described previously (*31*). For each cell, 500 surrogate position-tuning curves were prepared by randomly shifting the position-tuning curve within the represented paths. The PCC of cell *i* to event *e* was computed as follows:

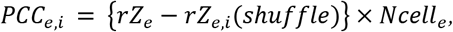

where *rZ*_*e,i*_*(shuffle)* is the averaged *rZ*_*e*_ obtained from the shuffled tuning curves of cell *i*, and *Ncell*_*e*_ is the number of activated place cells that encode the represented path in the event. For each cell, the PCCs were averaged over all events (Fig. 3C) or within each path (Supplementary Fig. 11).

#### The proportions of forward and reverse replays

Replay events for the start and goal areas were excluded from this analysis. Hierarchical Bayesian modeling was applied to estimate the posterior probability of the proportions of forward replays with the parameters estimated by Markov chain Monte Carlo (MCMC) methods. We assumed that the forward and backward replays in each trial emerged according to the binomial distribution as follows:

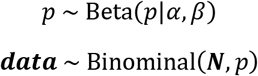

where **data** is a *T* × 1-dimensional vector containing the number of forward replays in each trial, ***N*** is a *T* × 1-dimensional vector containing the total number of replay events in each trial, the parameter *p* represents the binomial distribution, and the parameters α and β control the beta distribution and have initial values of 1. The parameter *p* was plotted as the proportion of forward replays in Fig. 3D and Supplementary Fig. 12E.

#### Trajectory events

A trajectory event was defined from a synchronous event, as previously described (*5, 34*). For each synchronous event, the posterior probability of the animal’s location *Pr(loc|s)* was computed by Bayesian decoding in which a 20-ms sliding window was moved every 10 ms as described above (*loc*: location, *s*: spikes). In the *Pr(loc|s)*, the maximum a posteriori probability (MAP) was computed for each path per time bin. Bins with MAPs greater than (5 × 1/*n*_*loc bin*_) were included in the following analyses, where *n*_*loc bin*_ is the total number of all location bins.

A candidate trajectory sequence was detected when the distances between all pairs of neighboring locations giving the MAP (allowing a 1-gap time bin) were less than 40 cm within the represented paths. The candidate trajectory sequences were then extended to the neighboring paths using the same criteria, and the sequence with the maximum length was adopted as a trajectory sequence. Trajectory sequences covering more than 4 time bins (60 ms) and a cumulative distance of more than 20 cm were included in the further analyses.

To assess the significance of a trajectory sequence, p-values were computed to represent the probability of detecting trajectory events from the 1,000 surrogate synchronous events with randomized position tuning curves. These surrogate synchronous events were created using the same procedure as described above in the analysis for *repZ*. Trajectory events were defined from the trajectory sequences where *p* < 0.05.

#### Proportion of trajectory events

Hierarchical Bayesian modeling was applied to estimate the posterior probability of the proportion of trajectory events per synchronous event with the parameters estimated by Markov chain Monte Carlo (MCMC) methods. We assumed that the trajectory events in each trial emerged according to a binomial distribution as follows:

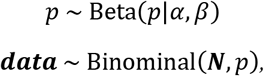

where **data** is a *T* × 1-dimensional vector containing the number of trajectory events in each trial, ***N*** is a *T* × 1-dimensional vector whose entries are the total number of synchronous events in each trial, the parameter *p* represents the binomial distribution, and the parameters α and β control the Beta distribution and have initial values of 1. The parameter *p* was plotted as the proportion of trajectory events in Fig. 3G and Supplementary Fig. 13B and 13C.

#### Trajectory-to-trajectory distance

The convergence of behavioral trajectories across trials in the presence of closed-loop stimulation was quantified in Fig. 4. Specifically, we analyzed the trajectories after the rat first took path start-S-C_2_ during the learning phase. A trajectory in the open field in each trial was converted to a character string (examples are shown in Supplementary Fig. 1F). A trajectory-to-trajectory distance was computed between two character strings in neighboring trials as a modified Levenshtein distance, with cost values considered as follows. A cost value for a string substitution was set to be (*distance*/1.13), where *distance* is the distance between the centers of two lattices involved in the substitution, and 1.13 is the distance (m) between lattices ‘a’ and ‘y’ representing the maximum lattice-to-lattice distance in the field (Supplementary Fig. 1E). The cost value for a deletion or insertion was 0.40/2, where 0.40 is the median of the normalized distance between all pairs of lattices.

When the trajectory-to-trajectory distance exceeded 2, the trial was considered to be a trial representing behavioral change. The percentage of trials with behavioral changes was computed for each rat, (Fig. 4F).

#### Statistical analysis

All the data were analyzed using MATLAB and Python. The data are presented as the mean ± standard error of the mean (SEM), dot plots with mean values, or box plots. Posterior probability distributions derived from the hierarchical Bayesian models with Markov chain Monte Carlo (MCMC) methods (*35*) are presented as violin plots or as lines showing the median values with 50% and 95% credible intervals.

Comparisons of one-sample data were analyzed using a one-sample *t*-test versus a constant value. Multiple group comparisons were performed by Tukey’s tests (Fig. 1R, 3B, and 4F) or by Mann-Whitney U tests followed by Bonferroni corrections (Fig. 1P). The null hypothesis was rejected at the *p* < 0.05 level. Posterior probability distributions of the estimated parameters with hierarchical Bayesian modeling were considered to be significantly different when their probability of overlap was less than 0.05 (Fig. 2H and 3F).

#### Data and software availability

The MClust software is available from A.D. Redish at http://redishlab.neuroscience.umn.edu/MClust/MClust.html. Reasonable requests for data and software will be fulfilled by the corresponding authors.

**Fig. S1.**
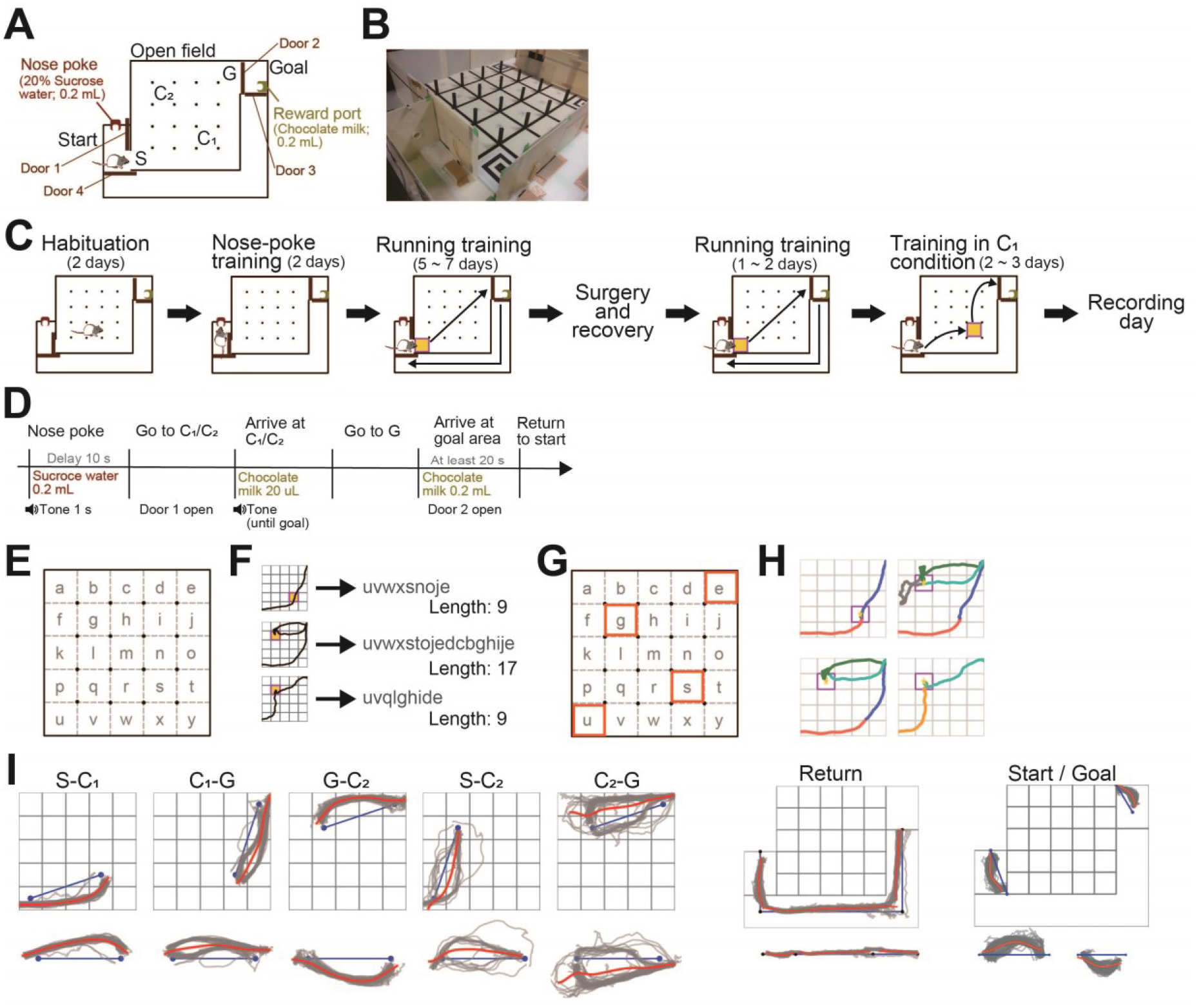
Behavioral apparatus for the spatial learning task and analysis of animal trajectories. (A) An overview of the area for the spatial learning task. “Nose poking” into the reward port in the start area for 1 s triggered the sound presentation and the dispensation of 25 × 8 **μ**l of 30% sucrose from the port. Next, door 1 was opened so that the rat could enter the open field and visit a checkpoint (C_1_ or C_2_) to obtain a 20 **μ**l chocolate milk reward (at one of the points C_1_ or C_2_). After reaching G, door 2 was opened so that the rat could enter the goal area. The reward port in the goal area then dispensed 200 **μ**l of chocolate milk reward. Finally, door 3 and door 4 were opened so that the rat could return to the start area via the return path. (B) A picture of the apparatus, viewed from the start area. (C) Timeline of training for the spatial learning task. (D) Time course of a single trial. (E) To analyze the trajectory patterns, the field was evenly divided into 5 × 5 lattices labeled with the letters “a-y”. The lattices u, e, s, and g correspond to S (after start), G (before goal), C_1_, and C_2_, respectively. (F) Three examples showing how the animals’ trajectories were analyzed. A trajectory in the field was converted into a character string (target string, left), and the total number of characters in of each string was counted as the string length. (G) The paths connecting the pairs of four specific lattices (u(S), e(G), s(C_1_), and g(C_2_)) were analyzed. The majority of segments were classified into S-C_1_, C_1_-G, G-C_2_, S-C_2_, and C_2_-G, which have a string length of 5. (H) Four representative rat trajectories are shown. A trajectory was divided into trajectory segments between the specific lattices, and each trajectory segment was classified into a path if the length of the segment was less than three plus the length of the path. Path segments that did not meet this criterion were classified as “other”. In the representative trajectories, each trajectory segment was colored based on the path (red, S-C_1_; blue, C_1_-G; green G-C_2_; orange, G-C_2_; cyan, C_2_-G). (I) For each trajectory segment, a straight line connecting the centers of the two lattices at both ends of the path type (blue line) was depicted for projection. The blue line was aligned so that its start point was set as the origin and moved onto the x-axis by an affine transformation (bottom panels). Individual rat trajectory segments (gray traces) were projected onto the same axis. All the trajectories were divided into 2-cm bins on the line, and an averaged trajectory (red traces) was computed on the line.

**Fig. S2.**
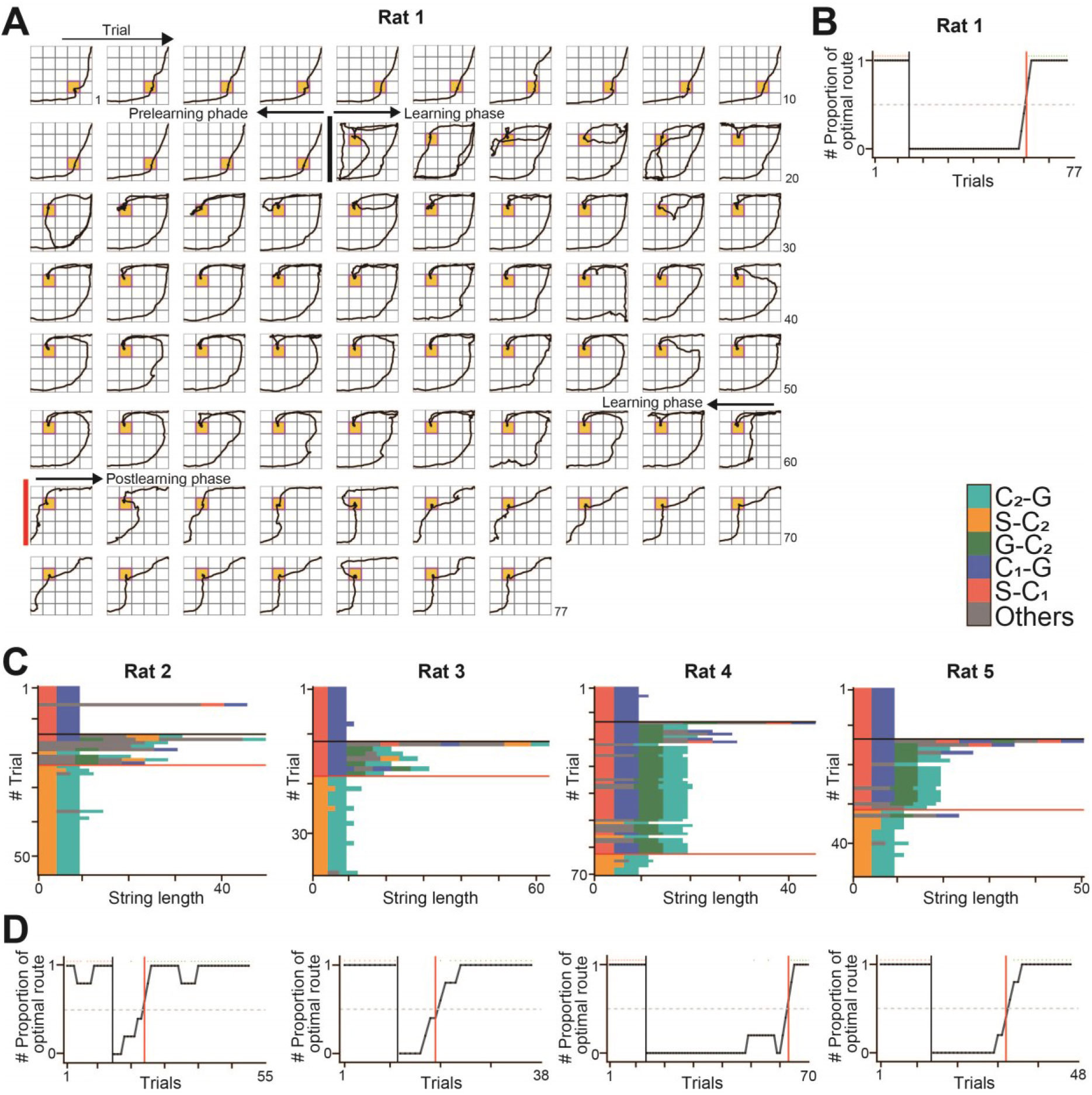
Behavioral results of all rats. (A) All the trajectories from a single rat (rat 1), corresponding with Fig. 1B. The black and red vertical lines indicate the reward replacement from C_1_ to C_2_ and the learning point, respectively. (B) A moving averaged learning curve, corresponding with A. (C) Changes in string length for each trial for the other four rats (rats 2-5). Data were plotted in the same way as in Fig. 1E. The black and red horizontal lines indicate the replacement and learning points, respectively. (D) Same as B but for the rats corresponding with C.

**Fig. S3.**
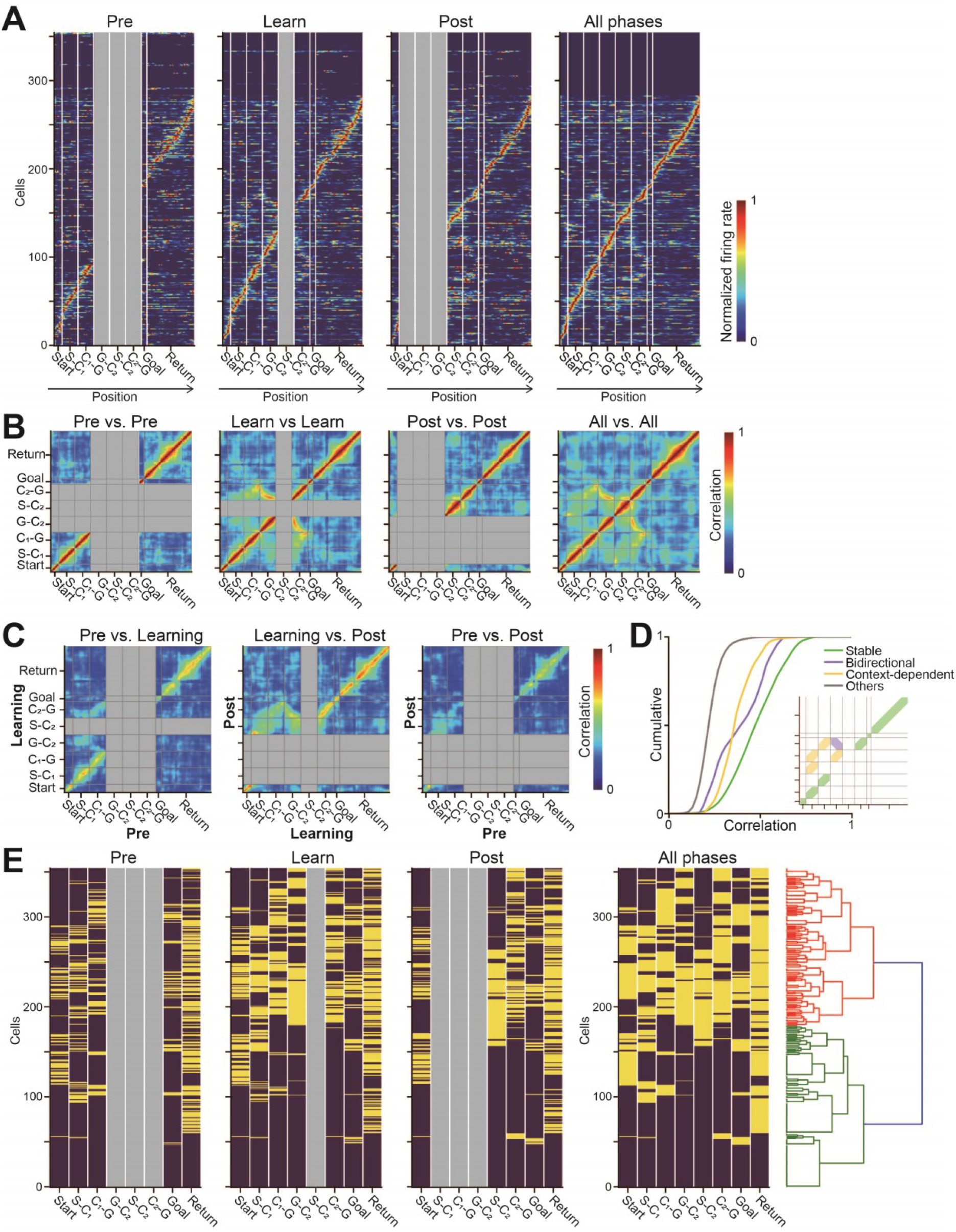
Spatial firing patterns of all recorded hippocampal cells. (A) Normalized spatial firing-rate distributions of 355 recorded hippocampal neurons from 5 rats, ordered by the locations of their place field center computed from all sessions. Locations not visited by the rat did not visit are shown in gray. The data are separately shown for the prelearning, learning, postlearning phases, and all phases combined. (B) Spatial correlation matrices of population vector pairs at all location bins. The locations not visited by the rat are shown in gray. (C) To assess the changes in spatial firing patterns associated with learning a new rewarded location, spatial cross-correlation matrices were constructed from population vector pairs at all the location bins. Overall, the spatial correlations at similar locations were higher than the spatial correlations at different locations, demonstrating stable spatial representation by place-cell ensembles. Higher density correlations were detected at the comparison path C_2_-G versus path G-C_2_ in the learning phase, implying the presence of place fields independent of running directions, termed “bidirectional fields”. Higher density correlations were also detected at S-C_1_ versus S-C_2_ and at C_1_-G versus C_2_-G, implying the presence of context-dependent fields relative to check points. (D) The cumulative distributions of correlation coefficients in specific regions (green, stable; yellow, context-dependent; purple, bidirectional) in the correlation matrices. The right bottom inset shows a schema depicting stable, context-dependent, and reverse fields (labeled by colors). (E) (Four left panels) Place-field locations of all recorded cells in individual phases (on, yellow; off, violet). The rightmost panel shows the hierarchical clustering of all recorded neurons based on their place-field locations.

**Fig. S4.**
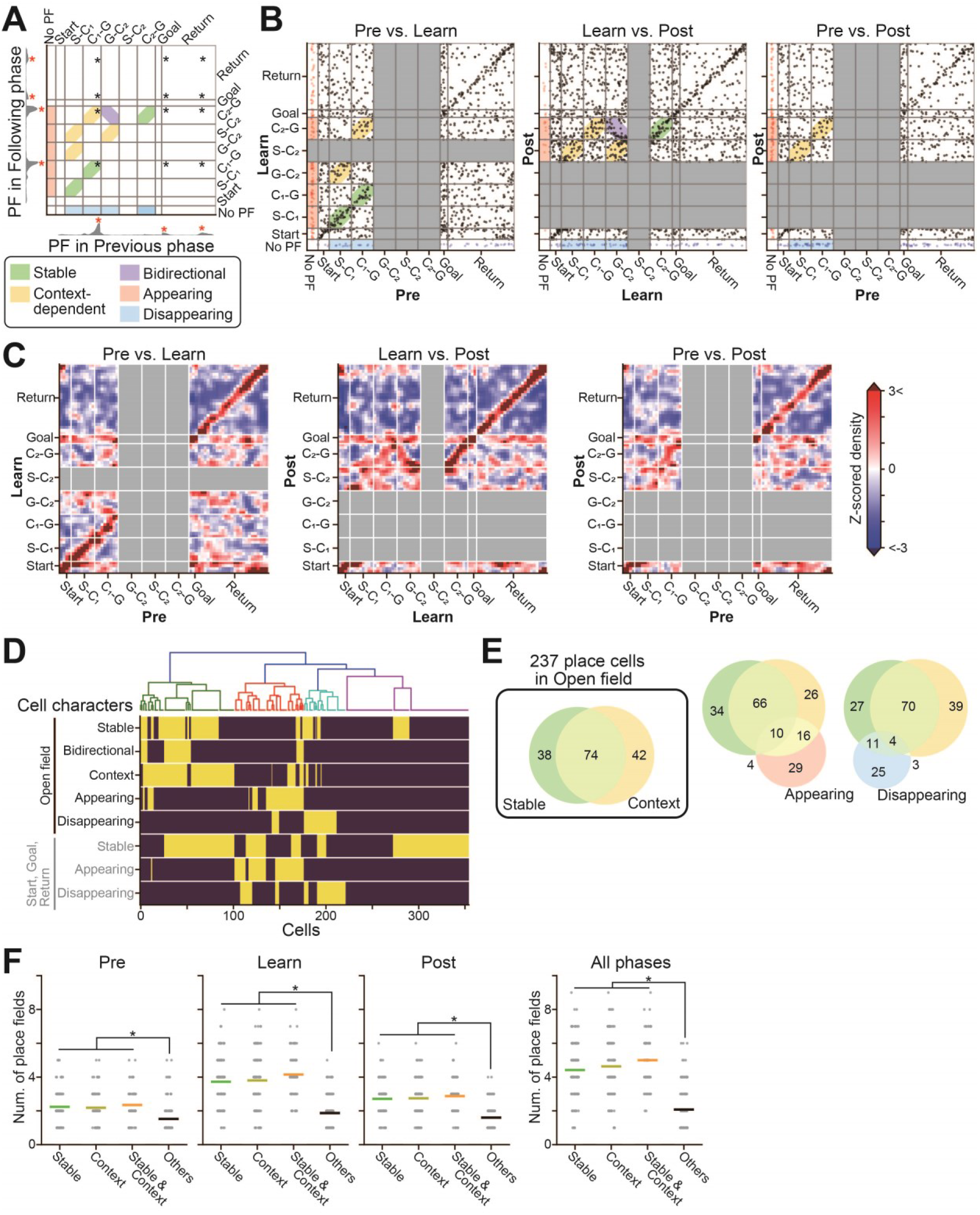
Learning-dependent changes in spatial maps. (A) This image is the same as the joint spatial map in Fig. 1K. Place fields (shown in red *) from a single place cell were defined based on spatial firing-rate distributions, and the locations of these place fields were plotted on the X- and Y-axes as black asterisks on a two-dimensional plain. (B) This image is the same as the joint spatial map in Fig. 1L but analyzed separately for all phase pairs. All the place fields from all cells were superimposed. The plots in the thin blue and red regions before the start area represent place fields that were not observed in one phase but emerged in the other phase. (C) Density plots of z-scores, constructed from B. In each bin, the density of the dots in B was z-scored based on the average and standard deviation of 1,000 surrogate datasets in which the locations of place fields at the latter phase were randomly shuffled without changing the total number of fields. Higher densities are visible, as shown in Supplementary Fig. 3B and 3C. (D) (top) Hierarchical clustering of all recorded neurons based on their field properties. (bottom) Summary of the field properties of all cells (on, yellow; off, violet). (E) Venn diagram showing the numbers of cells with individual field properties. (F) Comparisons of the number of place fields per place cell across cell types. Each gray dot represents a cell, and the colored lines represent the averages: *\*p* < 0.05, Mann-Whitney U test followed by Bonferroni correction.

**Fig. S5.**
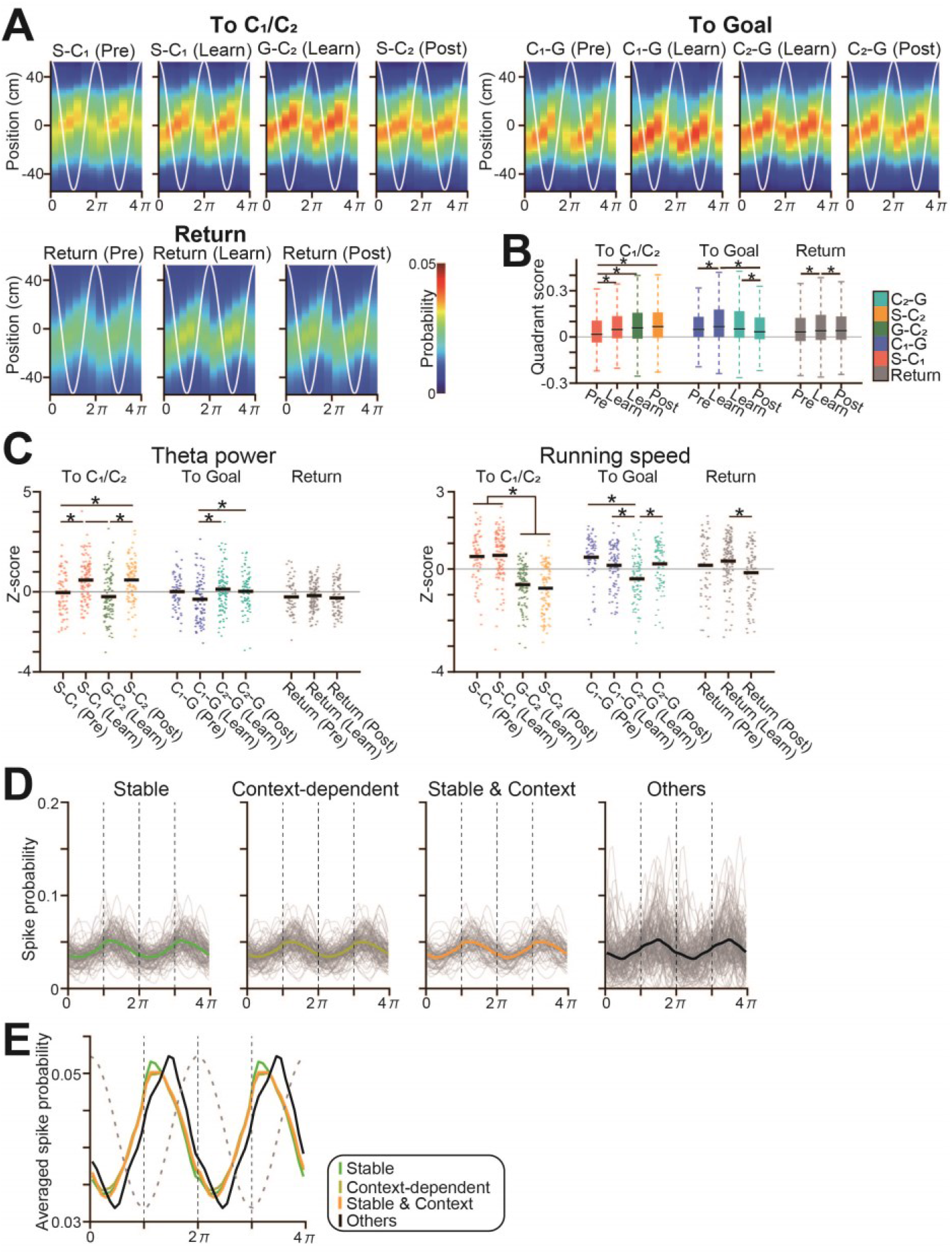
Learning-dependent and cell-type specific theta sequence. (A) Average posterior probabilities of animals’ positions while running on the path indicated above; the x-axis shows the phases of two theta cycles (white line) and the y-axis shows the positions relative to the current animal’s location. (B) The bar graphs shown in Fig. 1R are presented as boxplots: *\*p* < 0.05, Tukey’s test. (C) Theta power and running speed. Each dot represents a trial, and the black lines represent the averages: *\*p* < 0.05, Tukey’s test. (D) Cells were classified into stable, context-dependent, both stable and context-dependent, and other cells, based on Fig. 1L. The spike probabilities of individual pyramidal cells were plotted against the phase of theta oscillations. Each gray line represents one cell, and the colored lines represent the averages. (E) Superimposition of the average spike probabilities in each cell type corresponding with D.

**Fig. S6.**
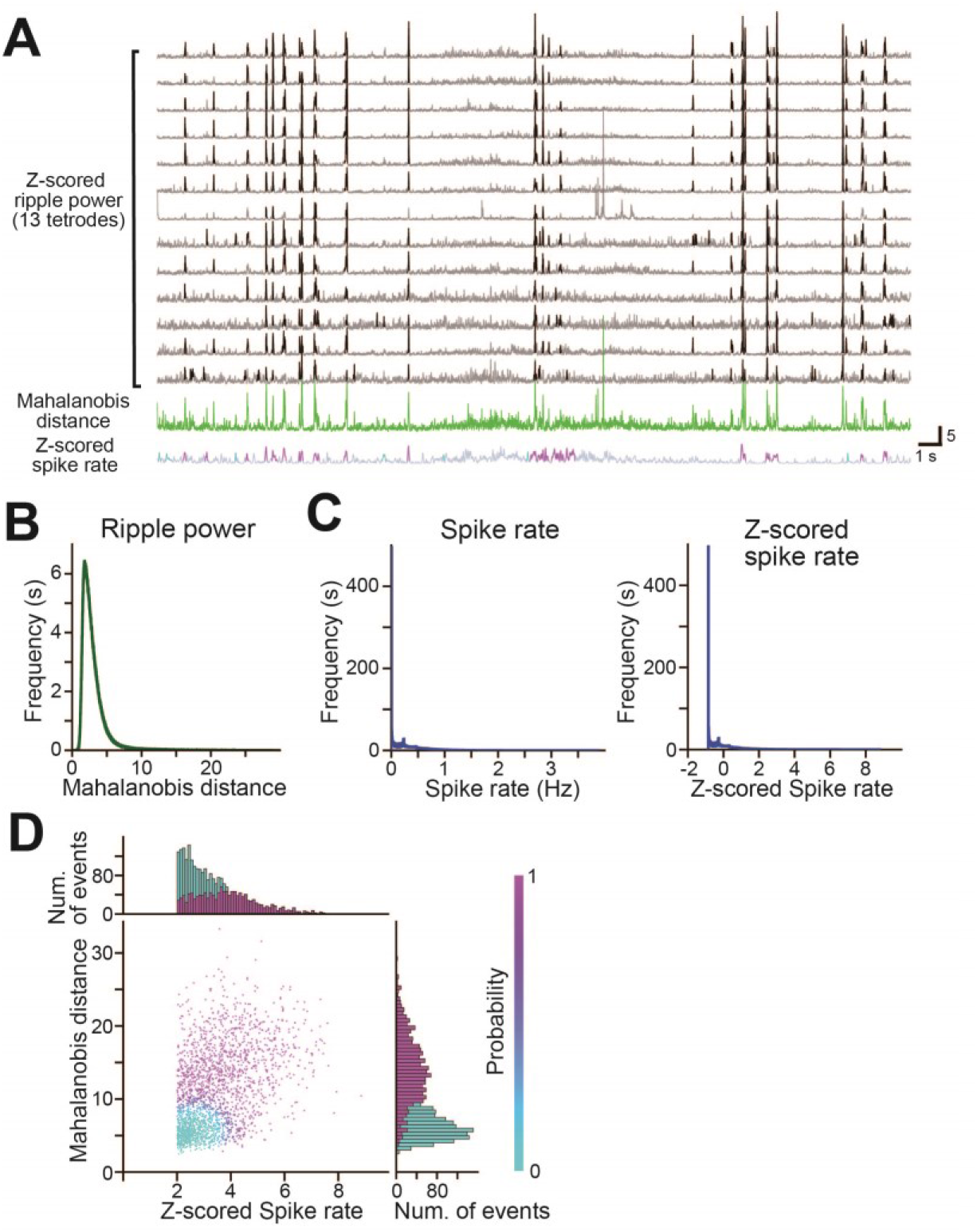
Detection of SWRs and synchronous events. (A) From top to bottom are the time changes in z-scored ripple power simultaneously recorded by thirteen tetrodes (upper gray traces), the corresponding Mahalanobis distance computed from the thirteen 150-250 Hz bandpass-filtered LFP traces as overall ripple power (green trace), and the z-scored spike rates of all recorded hippocampal cells (bin = 1 ms; sigma = 15 ms for Gaussian filter). Ripple power traces were computed from the 150-250 Hz bandpass-filtered LFP traces by the Hilbert transform (sigma = 4 ms for Gaussian filter), and the times at which ripples were detected in individual electrodes are colored in black. (B, C) The frequency distributions of the Mahalanobis distance (B) and the average spike rates of hippocampal cells: (C, left) (bin = 1 ms). The average spike rates are z-scored in the rightmost panel. (D) Relationship between the Mahalanobis distance and z-scored spike rates. Synchronous events were defined when the dots were included in the magenta cluster.

**Fig. S7.**
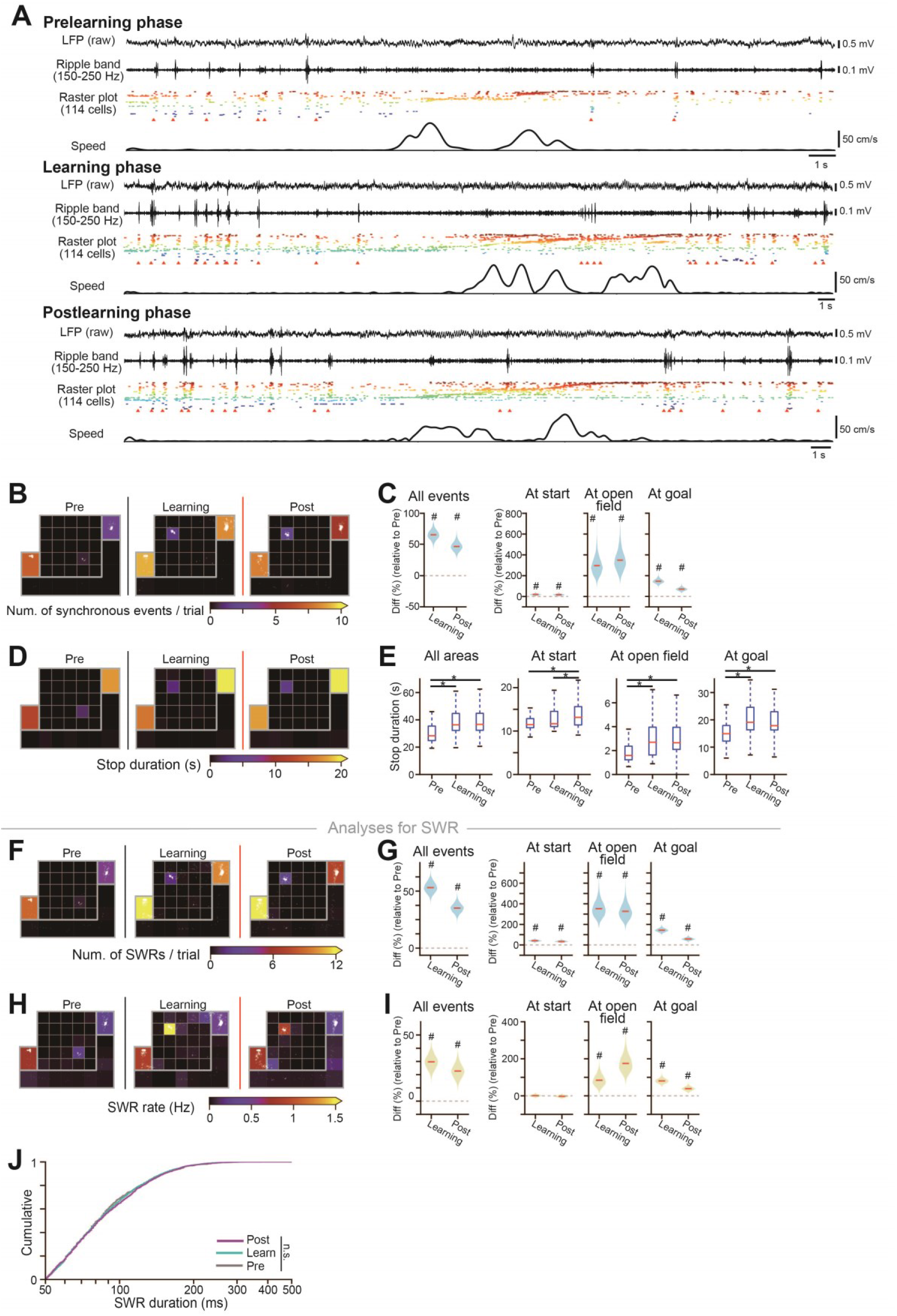
Learning-dependent changes in the numbers of synchronous events and SWRs. (A) From top to bottom are the original LFP, ripple band-filtered (150-250 Hz) LFP traces, a raster plot of the spike patterns of 114 neurons where the bottom arrowheads indicate synchronous events, and rat running speed during a single trial of each phase. Compared with the learning and postlearning phases, smaller numbers of synchronous events and SWRs are visible in the prelearning phase. (B) Pseudocolor maps of the average number of synchronous events during stop periods in each trial. The locations of synchronous events are indicated by superimposed white dots. (C) Same as Fig. 2C and 2D but analyzed for the absolute number of synchronous events relative to those in the prelearning phase. A pound sign (#) indicates no overlap between 0 and the 95% credible intervals computed from the posterior probability distribution by MCMC. (D) Pseudocolor maps of the average stop duration (moving speed less than 5 cm/s) in each trial. Comparisons of stop duration among the prelearning, learning, and postlearning phases. *\*p* < 0.05, Mann-Whitney U test followed by Bonferroni correction. (F, G) Same as B and C but analyzed for the number of SWRs. (H, I) Same as B and C but analyzed for the frequency of SWRs. (J) Cumulative distributions of SWR duration. No significant differences were found across the phases: *p* > 0.05, Mann-Whitney U test followed by Bonferroni correction.

**Fig. S8.**
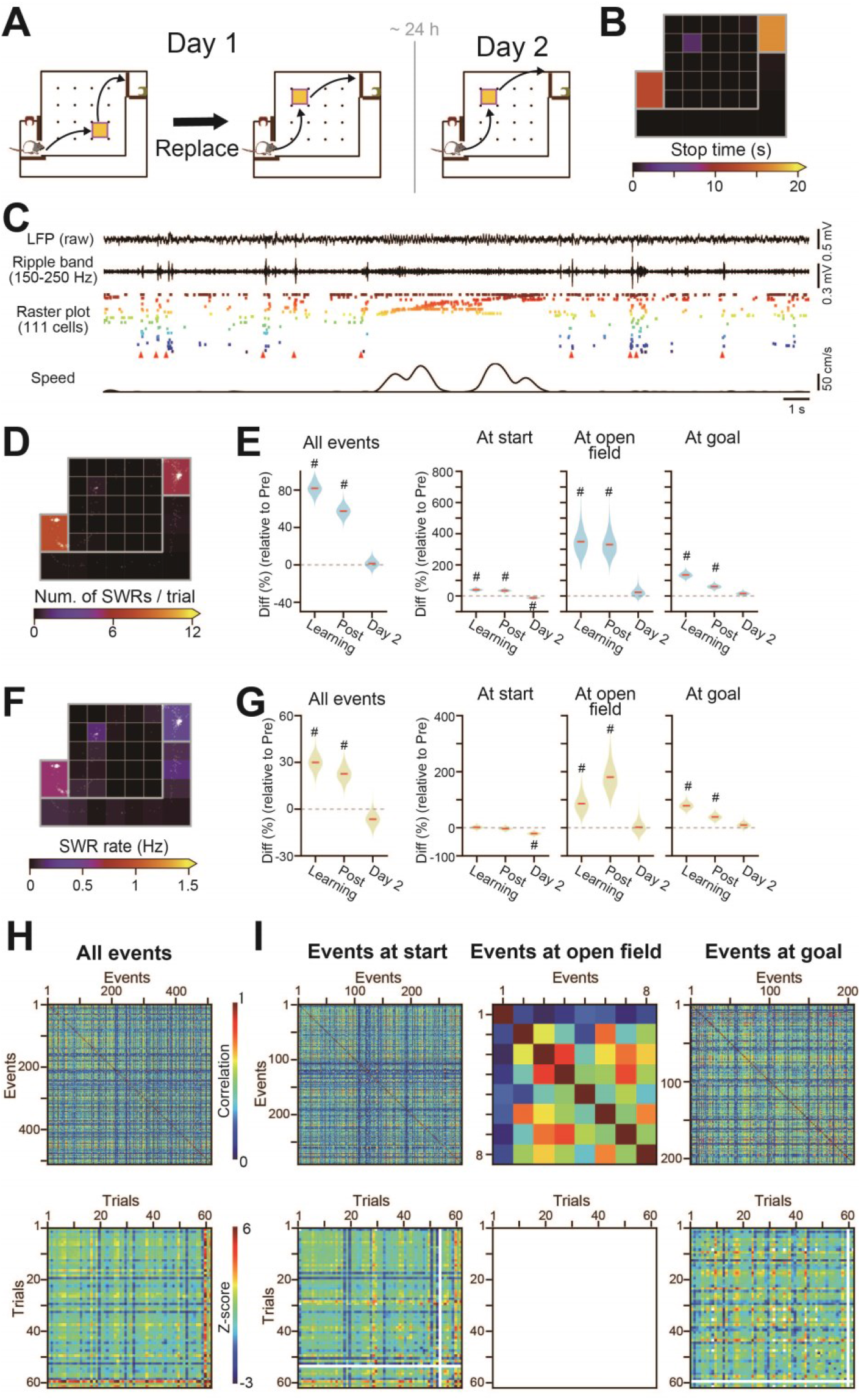
No increases in SWRs were observed under any learning conditions. (A) On the next day after the rat fully learned a checkpoint (C_2_), the same recording commenced without a change to the reward point (day 2), meaning that no further learning was induced. In this figure, all the analyses were performed from the data collected on day 2 unless otherwise specified. (B) A pseudocolor map of the average stop duration (moving speed less than 5 cm/s) in each trial on day 2. (C) From top to bottom are the original LFP, ripple band-filtered (150-250 Hz) LFP traces, a raster plot of spike patterns where the bottom arrowheads indicate synchronous events, and rat running speed from in a representative trial on day 2. Note that there are fewer synchronous events and SWRs compared with those in Fig. 1H. (D, F) Pseudocolor maps showing the number (D) and frequency (F) of SWRs per trial using the same color scales as in in Supplementary Fig. 7F and 7H. The locations of synchronous events are indicated by superimposed white dots. (E, G) The percentage of changes in the number and frequency of SWRs during the learning and postlearning phases on day 2 compared to the prelearning phase. Data from the prelearning, learning and postlearning phases are similar to those shown in Supplementary Fig. 7G and 7I for comparison. On day 2, no significant increases in SWRs were observed compared with the prelearning phase. A pound sign (#) indicates no overlap between 0 and the 95% credible intervals computed from posterior probability distribution by MCMC (*n* = 4-5 rats). (H) (top) An event-to-event correlation matrix of synchronous events on day 2. Detailed explanations are provided in Supplementary Fig. 9. (bottom) A trial-to-trial correlation matrix constructed from the event-to-event matrix in which the correlation coefficients are shown as z-scores computed from 1,000 surrogate datasets, showing no prominent changes in coefficients throughout the experiment, compared with those in Supplementary Fig. 9B-D. (I) Similar to H but separately analyzed for the individual areas. Trials with fewer than 3 synchronous events were not analyzed and are shown in white.

**Fig. S9.**
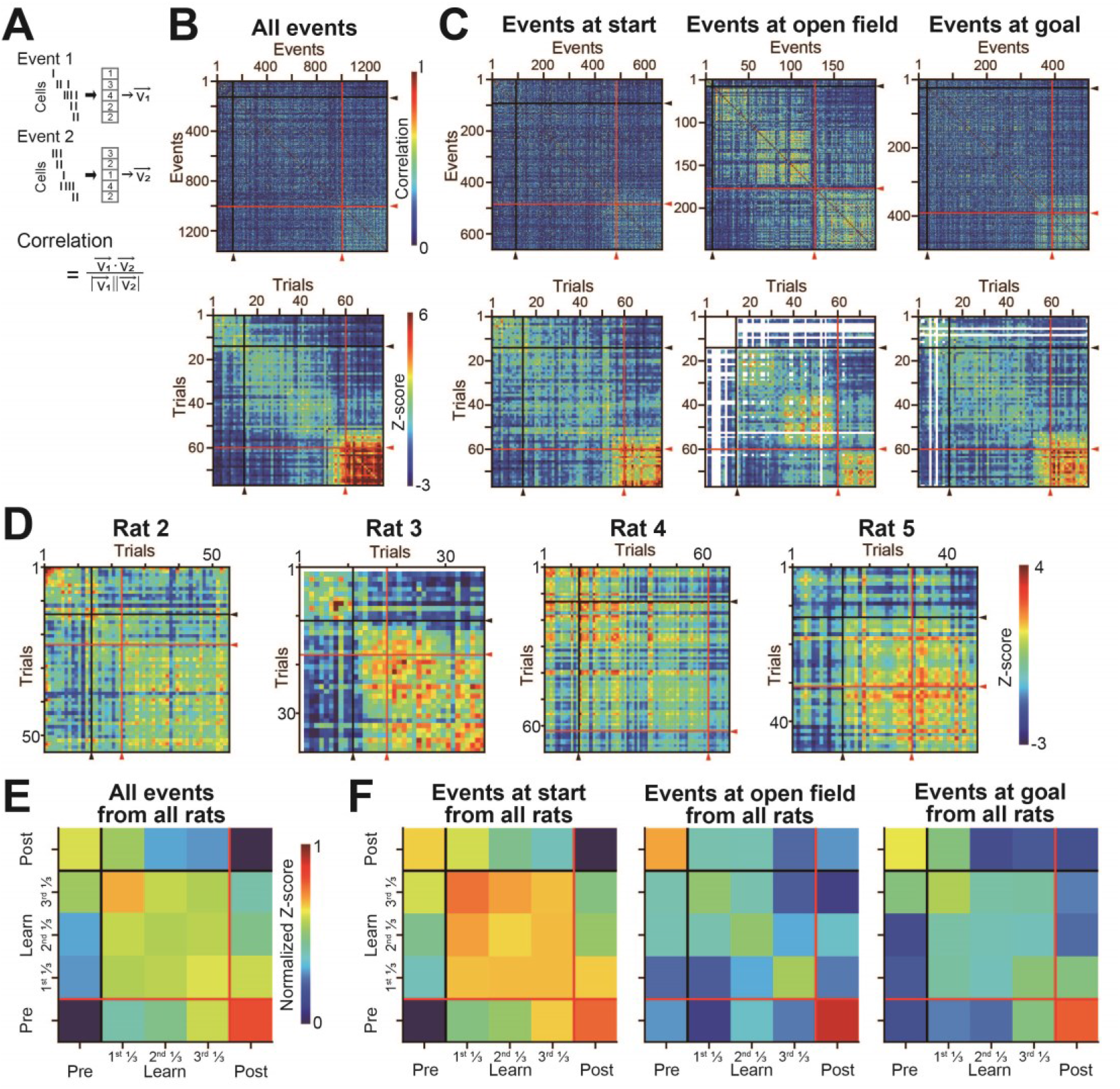
Correlation of spike patterns in synchronous events. (A) A population vector was constructed from the spike counts of all cells for each synchronous event. Correlation coefficients were computed from all pairs of population vectors and used to construct an event-to-event correlation matrix. (B) (top) An event-to-event correlation matrix of all synchronous events observed at all areas for one rat (rat 1). The black and red lines indicate the reward replacement from C_1_ to C_2_ and the learning point, respectively. (bottom) A trial-to-trial correlation matrix constructed from the event-to-event matrix by calculating the average of all the correlation coefficients included in each trial pair. Correlation coefficients are shown as z-scores based on 1,000 surrogate datasets. (C) Similar to B but separately analyzed for the individual areas. Trials with fewer than 3 synchronous events were not analyzed and are shown in white. Higher correlations are visible within each learning and postlearning phase compared with those within the prelearning phase. (D) Trial-to-trial correlation matrices in the other individual rats (rats 2-5). (E, F) Similar to the trial-to-trial correlation matrix, a phase-to-phase z-scored correlation matrix was constructed from the event-to-event matrix. A z-scored event-to-event matrix was first normalized so that the minimum and maximum values were 0 and 1, respectively, in each rat; then, the normalized matrices were averaged across all rats.

**Fig. S10.**
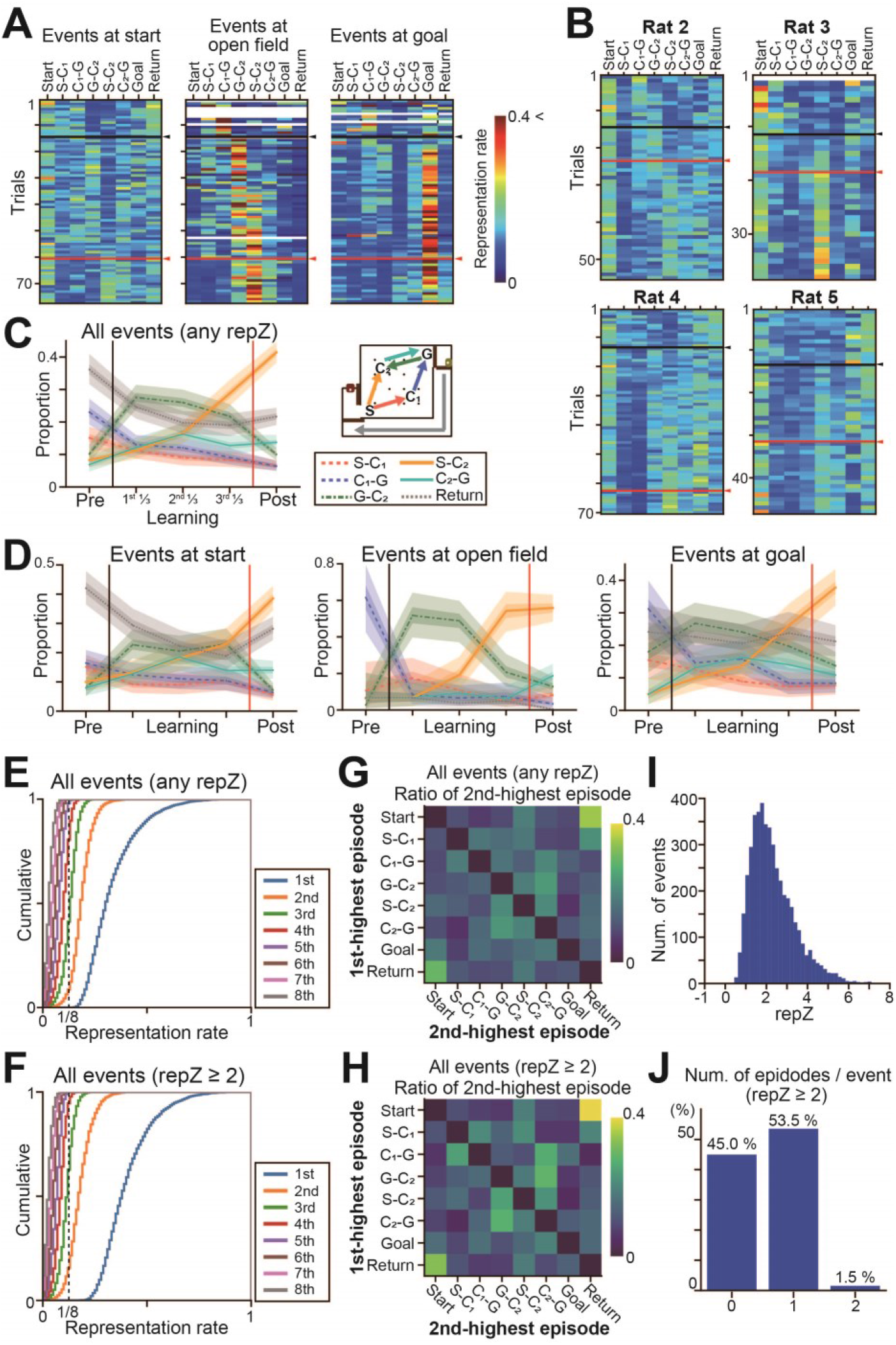
Represented paths by synchronous events. (A) Color-coded matrices showing changes in representation rates for each path by synchronous events, separately analyzed for individual areas (the same rat shown in Fig. 2F). The black and red lines indicate the reward replacement from C_1_ to C_2_ and the learning point, respectively. (B) Same as Fig. 2F but for the other individual rats (rats 2-5). (C) Same as Fig. 2G but for learning-related changes in the percentage of all synchronous events, irrespective of their *repZ*, representing individual paths. The thick and thin shaded areas indicate the 50% and 95% credible intervals, respectively. Similarly, pronounced increases in S-C_2_ are visible in the latter learning phases. (D) Same as C but separately plotted for the individual areas. (E, F) To analyze how the representation rates (*reprates*) of each synchronous event were biased for particular paths, z-scored representation rates (*repZ*) for all paths were computed for each synchronous event, and these *reprates* were ranked from highest (1st) to lowest (8th). For each rank, cumulative distributions were depicted from the *reprates* of all synchronous events (E) and synchronous events with *repZ* ≥ 2. (G, H) The probability of detecting synchronous events with the 1^st^ and 2^nd^ *reprate* at paths (or areas) indicated by the y-axis and x-axis, respectively. In H, a higher probability is visible in the comparison of neighboring paths (i.e., S-C_1_ versus C_1_-G and Start versus Return), implying the presence of joint replays of multiple episodes that are close in time and space. (I) The distribution of the 1^st^ *repZ* of all synchronous events. Synchronous events whose *repZ* ≥ 2 were considered to represent specific paths. (J) The number of represented paths identified from single synchronous events.

**Fig. S11.**
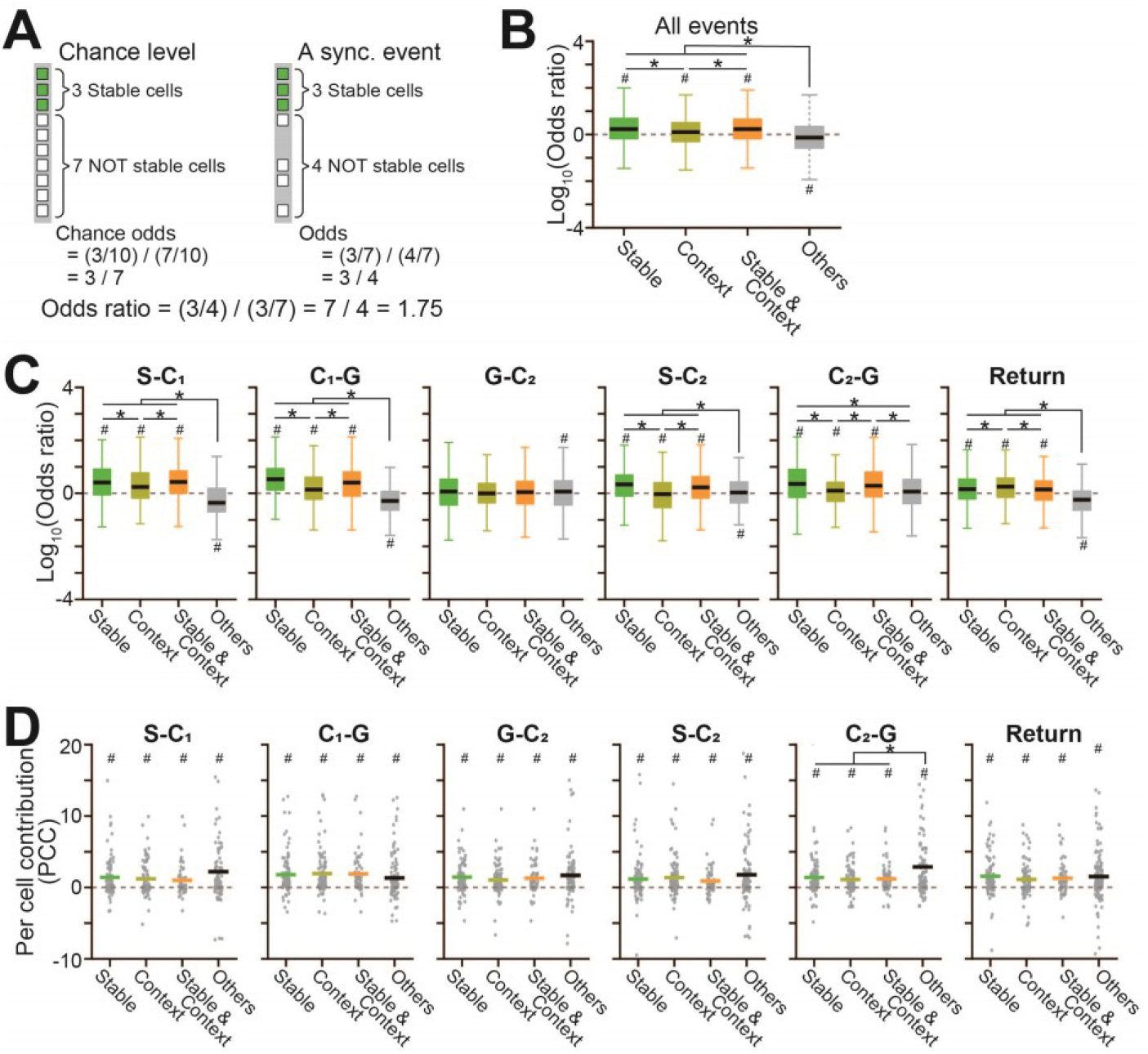
Cell participation in synchronous events and per cell contribution to replay events. (A) Schematic illustration of computing the odds ratios for a cell type. This illustration includes a total of 10 place cells with 3 stable cells (green boxes) and 7 other cells (white boxes). An example synchronous event includes 3 and 4 active cells. For this synchronous event, the odds were computed as the ratio of the percentage of active stable cells to total active cells (3/7) to the percentage of active other cells to total active cells (4/7). The odds at a chance level were computed as the ratio of the percentage of total stable cells to total cells (3/10) to the percentage of total other cells to total cells (7/10). To compute a chance level for each path, the active place cells in the path were analyzed. Finally, the odds ratio was computed as a ratio of the two odds as shown below. (B) Comparison of odds ratios across cell types: #*p* < 0.05, one-sample *t*-test versus 0; *\*p* < 0.05, Tukey’s test. (C) Same as B but separately analyzed for the individual paths. (D) Per cell contribution of individual cells to replay events, separately analyzed for the individual represented paths: #*p* < 0.05, one-sample *t*-test versus 0; *\*p* < 0.05, Tukey’s test.

**Fig. S12.**
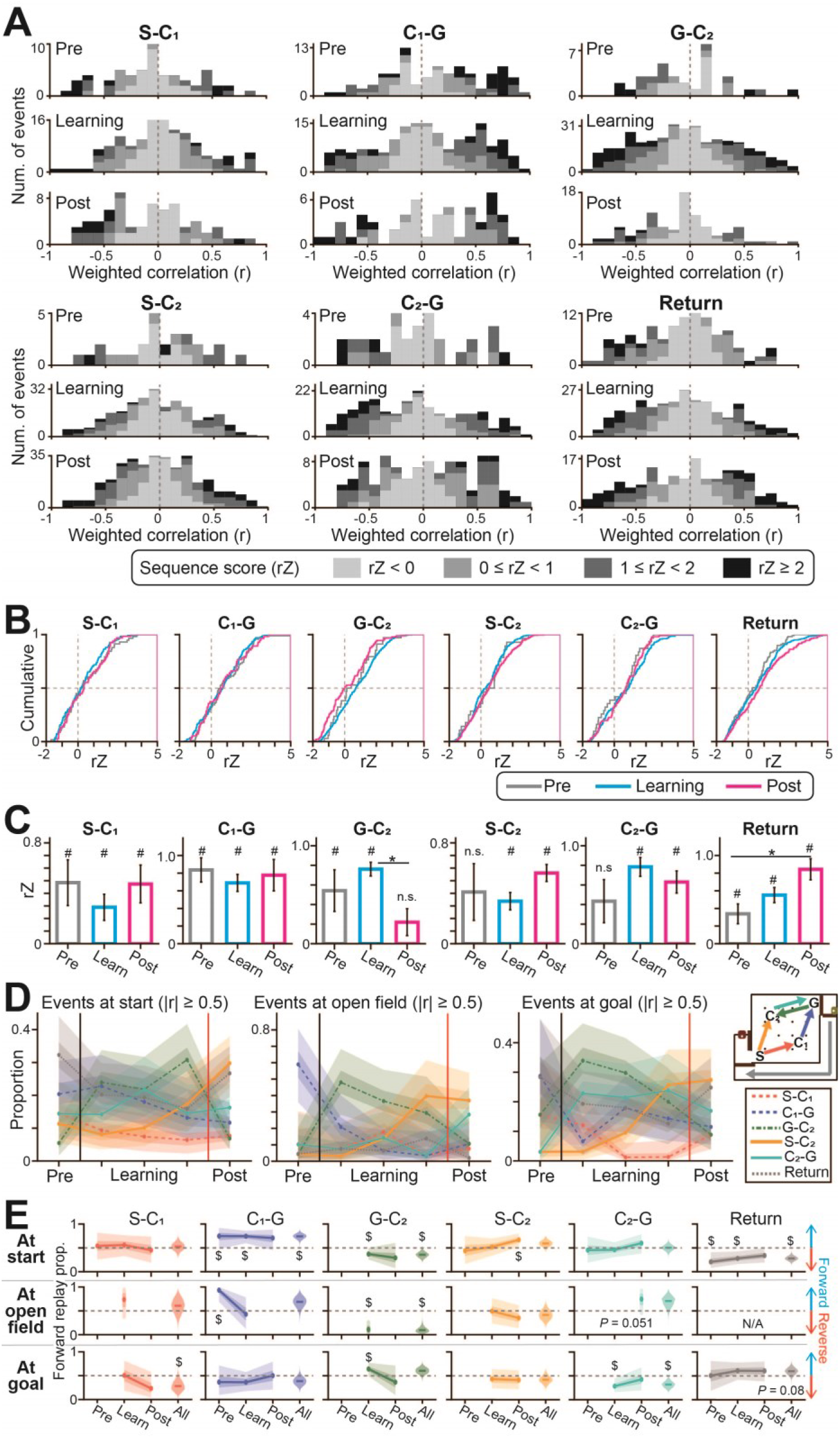
Sequential events representing individual paths. (A) Distributions of the weighted correlations (*r*) of synchronous events. The positive and negative *r* values represent forward and reverse directions, respectively. Each event is labeled in color depending on its sequence score (*rZ*). Synchronous events with an |*r*| above 0.5 were considered to be sequential events. (B) Cumulative distributions of the z-scored sequence scores (*rZ*) of synchronous events, separately analyzed for the individual represented paths. (C) The same data shown in B were averaged: *#p* < 0.05, one-sample t-test versus 0; \**p* < 0.05, Tukey’s test. (D) Same as Fig. 3F but separately analyzed for replay events in the individual areas. The thick and thin shaded areas indicate the 50% and 95% credible intervals, respectively. (E) The directionality of replay events separately analyzed for individual paths. The thick and thin shaded areas indicate the 50% and 95% credible intervals, respectively. A dollar ($) symbol indicates that the overlap in the distribution probability between 0.5 and the posterior probability distribution was less than 5%. Remarkably, the results showed that (1) replay events representing the return path observed at the start and goal areas were biased toward the reverse and forward directions, respectively, and (2) replay events representing paths G-C_2_ and C_2_-G observed in the open field (the majority of which emerged at C_2_ as shown in Fig. 2B) were biased toward the reverse and forward directions, respectively. These results demonstrate that forward and reverse replays preferentially emerge when their represented paths correspond with an animal’s behavioral episodes in the immediate future and past, respectively. Together with the results showing that reverse replays were more dominant than were forward replays in the open field (the majority of which emerged at C_2_) during the learning phase (Fig. 3D), path G-C_2_ is a main episode that is preferentially represented by reverse replays during the learning.

**Fig. S13.**
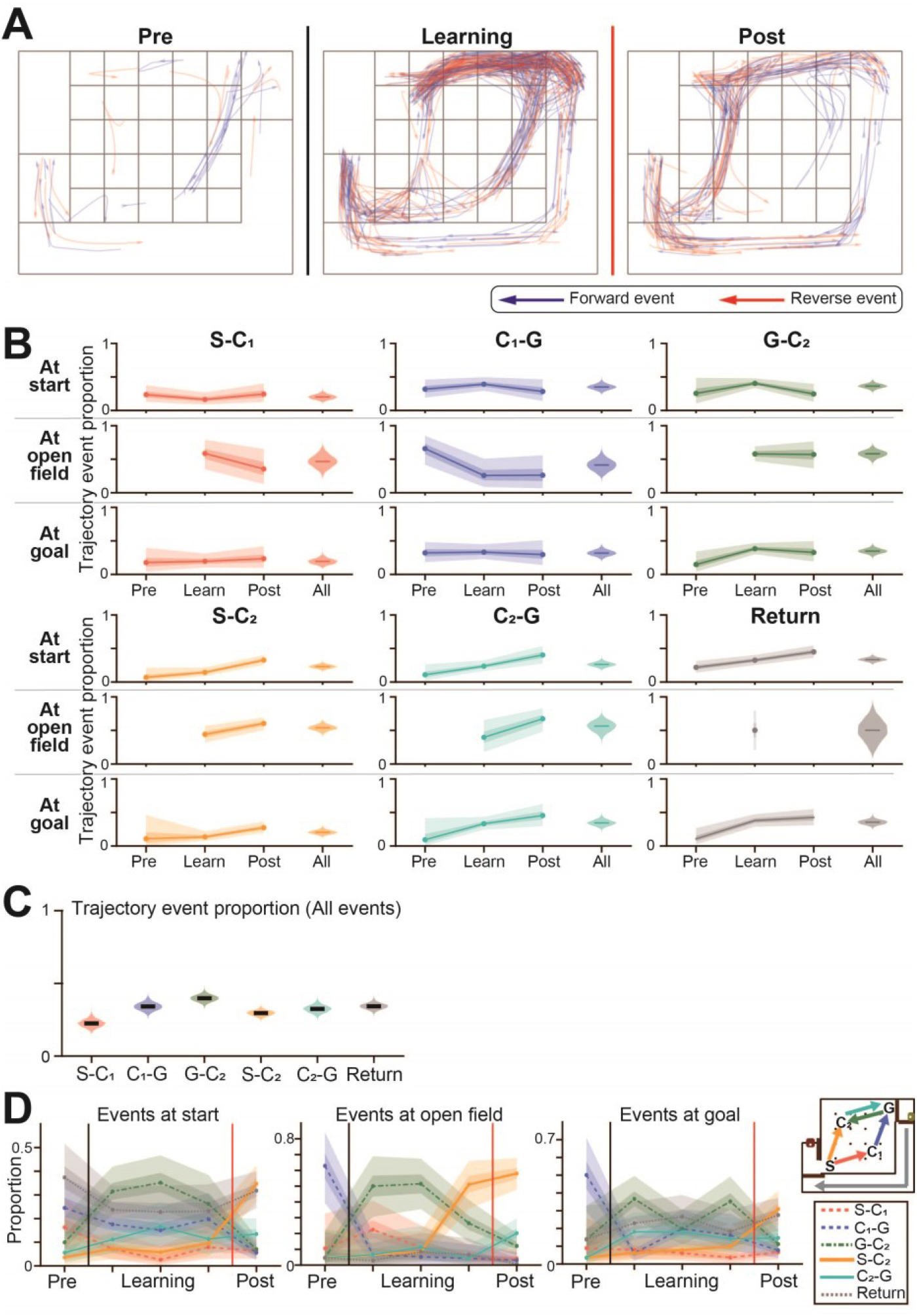
Representation by trajectory events. (A) Superimposition of the decoded trajectories from all trajectory events. The forward and reverse trajectory events are shown as blue and red lines, respectively. (B) The percentage of synchronous events to be assigned as trajectory events. The thick and thin shaded areas indicate the 50% and 95% credible intervals, respectively. (C) The same data shown in B were superimposed for comparison across paths. (D) The percentage of represented paths by trajectory events, separately analyzed for the individual areas. The thick and thin shaded areas indicate the 50% and 95% credible intervals, respectively.

**Fig. S14.**
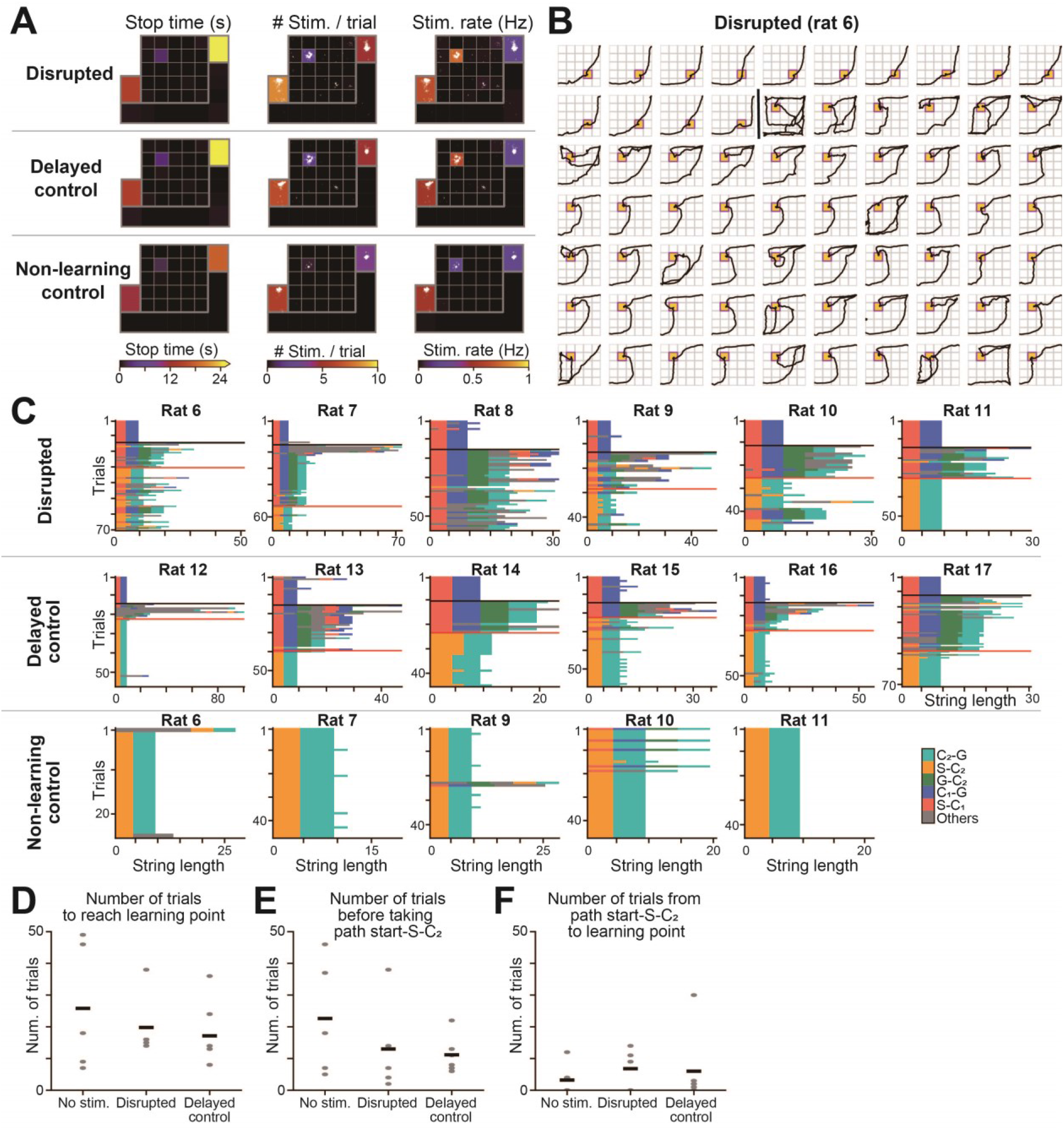
Effects of SWR inhibition on behavioral performance. (A) Pseudocolor maps of the average stop duration (moving speed less than 5 cm/s) and the number and frequency of feedback electrical stimulation applied per trial. Stimuli locations are indicated by superimposed white dots. (B) Representative running trajectories observed in a single rat with disrupted SWRs. The black vertical line indicates reward replacement. (C) Changes in string lengths for the other individual rats in the disrupted SWR, delayed control, and nonlearning control groups. The black and red horizontal lines indicate the reward replacement and the learning point, respectively. The data are colored depending on the paths taken by the rats. (D) The number of trials to reach the learning point in each group. Each dot represents a rat, and the black lines represent the averages. (E) The number of trials until the rats first took the path start-S-C_2_. (F) The number of trials from when rats first took the path start-S-C_2_ to the learning point.

**Fig. S15.**
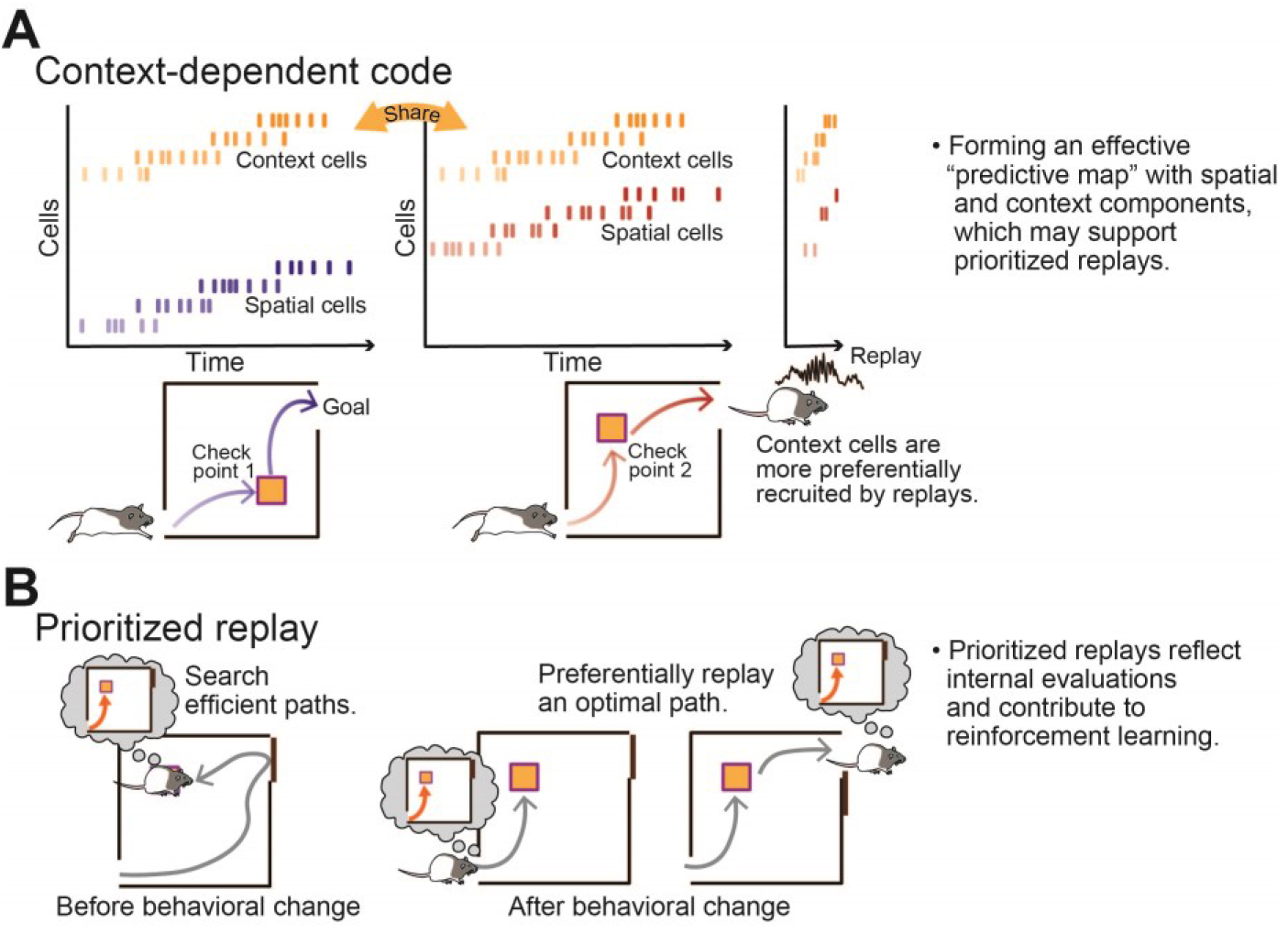
Schematic illustrations suggested from this study. (A) Context-dependent codes on a predictive map. Context-dependent cells, which encode task contexts (e.g. toward check points), are shared across learning phases and are preferentially recruited by hippocampal replays. The brain circuits form an effective predictive map with both spatial and context components, which may support memory processes and prioritized replays. (B) Prioritized replays on a predictive map. Hippocampal circuits preferentially replay salient episodes based on agent’s internal evaluations in a model in the brain. Such prioritized replays likely contribute to efficient learning.

## References

1. J. O’Keefe, L. Nadel, Oxford University Press, (1978).

2. K. L. Stachenfeld et al., Nat Neurosci 20, 1643–1653 (2017).

3. G. Buzsaki, Hippocampus 25, 1073–1188 (2015).

4. D. J. Foster, M. A. Wilson, Nature 440, 680–683 (2006).

5. B. E. Pfeiffer, D. J. Foster, Nature 497, 74–79 (2013).

6. S. Cheng, L. M. Frank, Neuron 57, 303–313 (2008).

7. J. O’Neill et al., Nat Neurosci 11, 209–215 (2008).

8. S. P. Jadhav et al., Science 336, 1454–1458 (2012).

9. L. Roux et al., Nat Neurosci 20, 845–853 (2017).

10. A. Johnson, A. D. Redish, Neural Netw 18, 1163–1171 (2005).

11. D. Hassabis et al., Neuron 95, 245–258 (2017).

12. V. Mnih et al., Nature 518, 529–533 (2015).

13. L. J. Lin, Machine Learning 8, 293–321 (1992).

14. A. C. Singer, L. M. Frank, Neuron 64, 910–921 (2009).

15. R. E. Ambrose et al., Neuron 91, 1124–1136 (2016).

16. M. G. Mattar, N. D. Daw, Nat Neurosci 21, 1609–1617 (2018).

17. T. Schaul et al., arXiv 1511.05952, (2015).

18. D. Aronov et al., Nature 543, 719–722 (2017).

19. Y. Liu et al., Cell 178, 640–652 e614 (2019).

20. Y. Aoki et al., Cell Rep 27, 1516–1527 e1515 (2019).

21. J. L. Gauthier, D. W. Tank, Neuron 99, 179–193 e177 (2018).

22. A. M. Wikenheiser, A. D. Redish, Nat Neurosci 18, 289–294 (2015).

23. C. Drieu et al., Science 362, 675–679 (2018).

24. Z. Brzosko et al., Elife 6, (2017).

25. K. J. Miller et al., Nat Neurosci 20, 1269–1276 (2017).

26. M. Botvinick et al., Trends Cogn Sci 23, 408–422 (2019).

27. T. E. J. Behrens et al., Neuron 100, 490–509 (2018).

28. A. Banino et al., Nature 557, 429–433 (2018).

## References

1. J. O’Keefe, L. Nadel, The Hippocampus as a Cognitive Map. Oxford University Press, (1978).

2. K. L. Stachenfeld, M. M. Botvinick, S. J. Gershman, The hippocampus as a predictive map. Nat Neurosci 20, 1643–1653 (2017). doi:10.1038/nn.4650.

3. G. Buzsaki, Hippocampal sharp wave-ripple: A cognitive biomarker for episodic memory and planning. Hippocampus 25, 1073–1188 (2015). doi:10.1002/hipo.22488.

4. D. J. Foster, M. A. Wilson, Reverse replay of behavioural sequences in hippocampal place cells during the awake state. Nature 440, 680–683 (2006). doi:10.1038/nature04587.

5. B. E. Pfeiffer, D. J. Foster, Hippocampal place-cell sequences depict future paths to remembered goals. Nature 497, 74–79 (2013). doi:10.1038/nature12112.

6. S. Cheng, L. M. Frank, New experiences enhance coordinated neural activity in the hippocampus. Neuron 57, 303–313 (2008). doi:10.1016/j.neuron.2007.11.035.

7. J. O’Neill, T. J. Senior, K. Allen, J. R. Huxter, J. Csicsvari, Reactivation of experience-dependent cell assembly patterns in the hippocampus. Nat Neurosci 11, 209–215 (2008). doi:10.1038/nn2037.

8. S. P. Jadhav, C. Kemere, P. W. German, L. M. Frank, Awake hippocampal sharp-wave ripples support spatial memory. Science 336, 1454–1458 (2012). doi:10.1126/science.1217230.

9. L. Roux, B. Hu, R. Eichler, E. Stark, G. Buzsaki, Sharp wave ripples during learning stabilize the hippocampal spatial map. Nat Neurosci 20, 845–853 (2017). doi:10.1038/nn.4543.

10. A. Johnson, A. D. Redish, Hippocampal replay contributes to within session learning in a temporal difference reinforcement learning model. Neural Netw 18, 1163–1171 (2005). doi:10.1016/j.neunet.2005.08.009.

11. D. Hassabis, D. Kumaran, C. Summerfield, M. Botvinick, Neuroscience-Inspired Artificial Intelligence. Neuron 95, 245–258 (2017). doi:10.1016/j.neuron.2017.06.011.

12. V. Mnih, K. Kavukcuoglu, D. Silver, A. A. Rusu, J. Veness, M. G. Bellemare, A. Graves, M. Riedmiller, A. K. Fidjeland, G. Ostrovski, S. Petersen, C. Beattie, A. Sadik, I. Antonoglou, H. King, D. Kumaran, D. Wierstra, S. Legg, D. Hassabis, Human-level control through deep reinforcement learning. Nature 518, 529–533 (2015). doi:10.1038/nature14236.

13. L. J. Lin, Self-Improving Reactive Agents Based on Reinforcement Learning, Planning and Teaching. Machine Learning 8, 293–321 (1992). doi:Doi 10.1023/A:1022628806385.

14. A. C. Singer, L. M. Frank, Rewarded outcomes enhance reactivation of experience in the hippocampus. Neuron 64, 910–921 (2009). doi:10.1016/j.neuron.2009.11.016.

15. R. E. Ambrose, B. E. Pfeiffer, D. J. Foster, Reverse Replay of Hippocampal Place Cells Is Uniquely Modulated by Changing Reward. Neuron 91, 1124–1136 (2016). doi:10.1016/j.neuron.2016.07.047.

16. M. G. Mattar, N. D. Daw, Prioritized memory access explains planning and hippocampal replay. Nat Neurosci 21, 1609–1617 (2018). doi:10.1038/s41593-018-0232-z.

17. T. Schaul, J. Quan, I. Antonoglou, D. Silver, Prioritized experience replay. arXiv 1511.05952, (2015).

18. D. Aronov, R. Nevers, D. W. Tank, Mapping of a non-spatial dimension by the hippocampal-entorhinal circuit. Nature 543, 719–722 (2017). doi:10.1038/nature21692.

19. Y. Liu, R. J. Dolan, Z. Kurth-Nelson, T. E. J. Behrens, Human Replay Spontaneously Reorganizes Experience. Cell 178, 640–652 e614 (2019). doi:10.1016/j.cell.2019.06.012.

20. Y. Aoki, H. Igata, Y. Ikegaya, T. Sasaki, The Integration of Goal-Directed Signals onto Spatial Maps of Hippocampal Place Cells. Cell Rep 27, 1516–1527 e1515 (2019). doi:10.1016/j.celrep.2019.04.002.

21. J. L. Gauthier, D. W. Tank, A Dedicated Population for Reward Coding in the Hippocampus. Neuron 99, 179–193 e177 (2018). doi:10.1016/j.neuron.2018.06.008.

22. A. M. Wikenheiser, A. D. Redish, Hippocampal theta sequences reflect current goals. Nat Neurosci 18, 289–294 (2015). doi:10.1038/nn.3909.

23. C. Drieu, R. Todorova, M. Zugaro, Nested sequences of hippocampal assemblies during behavior support subsequent sleep replay. Science 362, 675–679 (2018). doi:10.1126/science.aat2952.

24. Z. Brzosko, S. Zannone, W. Schultz, C. Clopath, O. Paulsen, Sequential neuromodulation of Hebbian plasticity offers mechanism for effective reward-based navigation. Elife 6, (2017). doi:10.7554/eLife.27756.

25. K. J. Miller, M. M. Botvinick, C. D. Brody, Dorsal hippocampus contributes to model-based planning. Nat Neurosci 20, 1269–1276 (2017). doi:10.1038/nn.4613.

26. M. Botvinick, S. Ritter, J. X. Wang, Z. Kurth-Nelson, C. Blundell, D. Hassabis, Reinforcement Learning, Fast and Slow. Trends Cogn Sci 23, 408–422 (2019). doi:10.1016/j.tics.2019.02.006.

27. T. E. J. Behrens, T. H. Muller, J. C. R. Whittington, S. Mark, A. B. Baram, K. L. Stachenfeld, Z. Kurth-Nelson, What Is a Cognitive Map? Organizing Knowledge for Flexible Behavior. Neuron 100, 490–509 (2018). doi:10.1016/j.neuron.2018.10.002.

28. A. Banino, C. Barry, B. Uria, C. Blundell, T. Lillicrap, P. Mirowski, A. Pritzel, M. J. Chadwick, T. Degris, J. Modayil, G. Wayne, H. Soyer, F. Viola, B. Zhang, R. Goroshin, N. Rabinowitz, R. Pascanu, C. Beattie, S. Petersen, A. Sadik, S. Gaffney, H. King, K. Kavukcuoglu, D. Hassabis, R. Hadsell, D. Kumaran, Vector-based navigation using grid-like representations in artificial agents. Nature 557, 429–433 (2018). doi:10.1038/s41586-018-0102-6.

29. A. D. Redish, MClust 3.5, Free-Ware Spike Sorting (University of Minnesota, Minneapolis). Available at http://redishlab.neuroscience.umn.edu/MClust/MClust.html., (2009).

30. N. Schmitzer-Torbert, J. Jackson, D. Henze, K. Harris, A. D. Redish, Quantitative measures of cluster quality for use in extracellular recordings. Neuroscience 131, 1–11 (2005). doi:10.1016/j.neuroscience.2004.09.066.

31. A. D. Grosmark, G. Buzsaki, Diversity in neural firing dynamics supports both rigid and learned hippocampal sequences. Science 351, 1440–1443 (2016). doi:10.1126/science.aad1935.

32. T. Feng, D. Silva, D. J. Foster, Dissociation between the experience-dependent development of hippocampal theta sequences and single-trial phase precession. J Neurosci 35, 4890–4902 (2015). doi:10.1523/JNEUROSCI.2614-14.2015.

33. X. Wu, D. J. Foster, Hippocampal replay captures the unique topological structure of a novel environment. J Neurosci 34, 6459–6469 (2014). doi:10.1523/JNEUROSCI.3414-13.2014.

34. A. A. Carey, Y. Tanaka, M. A. A. van der Meer, Reward revaluation biases hippocampal replay content away from the preferred outcome. Nat Neurosci 22, 1450–1459 (2019). doi:10.1038/s41593-019-0464-6.

35. A. Gelman, J. B. Carlin, H. S. Stern, D. B. Duncon, A. Vehtari, D. B. RUbin, Bayesian Data Analysis, Third Edition (CRC Press, 2013), (2013).

